# FHOD3 and DIAPH3 control cell migration and differentially shift the balance of parallel and perpendicular stress fibers

**DOI:** 10.64898/2026.02.13.703290

**Authors:** Fred Rogers Namanda, Azarnoosh Foroozandehfar, Ian C. Schneider

## Abstract

Cell morphology, dictated by the filamentous actin (F-actin) cytoskeleton, is fundamental to cell migration during wound healing and cancer metastasis. Cell morphology is shaped by the extracellular matrix (ECM), which provides mechanical cues in the form of ECM stiffness. These mechanical cues regulate the assembly of the F-actin cytoskeleton which in turn controls cell morphology and cell migration. Formins are key regulators of linear F-actin, assembling it into stress fibers, yet the specific roles of individual formins in controlling distinct stress fiber subpopulations to control cell morphology and migration remain poorly defined. Here, we characterize formin expression across different cell types and leverage the inherent expression and cell morphology differences to identify FHOD3 and DIAPH3 as strongly correlated with cell elongation. We demonstrate that these formins regulate complementary but distinct stress fiber networks. In contractile, but less motile cells, FHOD3 knock-down shifts the balance towards stress fibers oriented perpendicular to the long axis of the cell. In contrast, DIAPH3 knock-down shifts the balance towards stress fibers oriented parallel to the long axis of the cell. However, in less contractile and highly motile cells, knockdown of either formin significantly impairs cell migration speed, suggesting both F-actin fiber networks are necessary for cell migration. Our work establishes a model where FHOD3 and DIAPH3 function through non-overlapping mechanisms to control the F-actin architecture that governs cell shape and motility.

## 1. Introduction

Cell migration is crucial for physiological and pathological events, including embryonic development, immune response, tissue repair, and cancer metastasis. Cell migration depends on precise control over actin dynamics, which are physically linked to the extracellular matrix (ECM) (Alonso-Matilla et al., 2025; Conboy et al., 2024; Doyle et al., 2022). Migrating cells continuously sense and respond to these physical cues such as ECM alignment and topography (Foroozandehfar et al., 2025), tension (Alisafaei et al., 2025) and stiffness (Yang et al., 2025). These physical inputs induce particular morphologies in cells that allow them to effectively migrate or remodel the local ECM (Duggan and Petrie, 2025). Substrate stiffness in particular is a critical mechanical cue that governs cell morphology, cell migration and cytoskeletal organization (Allan and Chaudhuri, 2025; Yeung et al., 2005). Cells exhibit enhanced spreading, a polarized morphology and larger traction forces on more rigid substrates, while on softer substrates, they exhibit a rounded cell shape and reduced traction forces (Califano and Reinhart-King, 2010; Tee et al., 2011). While this is the general trend, cells can also spread in stiffness-independent ways under certain ECM conditions (Conway et al., 2023), resulting in a traction maximum at intermediate stiffnesses (Bangasser et al., 2017). Additionally, different cell types respond distinctly to substrate stiffness (Yu et al., 2021) at the level of morphology (Yeung et al., 2005) or migration (Bangasser et al., 2017). Cells sense stiffness through integrin-mediated adhesion and actomyosin contractility, enabling mechanotransduction pathways to alter morphology and motility (Discher et al., 2005). Actomyosin contractility is enabled through the assembly of F-actin structures within the cell. Several types of contractile F-actin structures are built including network F-actin composed of loosely organized linear F-actin fibers in the lamella and linear F-actin structures frequently called stress fibers (Weißenbruch and Mayor, 2024). These contractile fibers include transverse arcs, dorsal stress fibers and ventral stress fibers (Hotulainen and Lappalainen, 2006; Kassianidou and Kumar, 2015; Maninová and Vomastek, 2016; Vallenius, 2013). All can be embedded in a contractile network (Vignaud et al., 2021). Transverse arcs run parallel with the protruding cell edge and perpendicular to the long axis of the cell and terminate in focal adhesions, generating traction perpendicular to cell movement. Dorsal stress fibers are parallel with the long axis of the cells and link focal adhesions at the front of the cell to the transverse arcs. Ventral stress fibers are also parallel with the long axis of the cells and connect focal adhesions at the front and rear of the cell. Each population is differentially regulated by myosin (Kuragano et al., 2018) and exerts tension differently within the cells (Lee et al., 2018). Furthermore, parallel and perpendicular F-actin structures respond differently to strains (Amiri et al., 2023). Actin-nucleation and elongation proteins play a specialized role in assembling different F-actin stress fibers. One important family of F-actin nucleating and elongating proteins, which induce the assembly and formation of stress fibers are formins (Lehtimäki et al., 2021; Schulze et al., 2014; Tojkander et al., 2011).

Formins are a highly conserved family of actin elongating proteins (Breitsprecher and Goode, 2013) found throughout eukaryotes (Ahangar and Cowin, 2022). Literature has indicated that disruption of formins, with pathological mutations, is linked to human disease such as kidney disease, peripheral neuropathies, intellectual disability, hearing loss, and cardiomyopathy and alterations in expression lead to cancer metastasis. Often their mis-regulation is linked to diseases associated with cytoskeletal regulation and migration such as cancer metastasis (Labat-de-Hoz and Alonso, 2021). They contain a conserved core of Formin Homology 1 and 2 (FH1 and FH2) domains, where the FH2 domain directly nucleates new actin filaments and remains bound to the growing barbed end (Courtemanche, 2018; Pruyne et al., 2002), while the adjacent FH1 domain recruits profilin-actin to drive processive elongation (Paul and Pollard, 2008). Their unique ability to processively elongate linear actin filaments is critical for the formation of F-actin structures that regulate cell shape and migration (Heuser et al., 2020; Scholz et al., 2024; Sigler et al., 2024) through lamellipodia, stress fibers and other F-actin structures (Schirenbeck et al., 2005; Svitkina et al., 2003). Formins stabilize F-actin by protecting barbed ends from capping proteins and by bundling filaments into larger structures, which enhance mechanical stability and modulate contractility (Goode and Eck, 2007; Tojkander et al., 2012). FMNL1-3 enable membrane protrusion and force generation by the lamellipodium, essential processes for cell movement (Kage et al., 2017). DIAPH1-3 polymerize stress fibers out of focal adhesions (Hotulainen and Lappalainen, 2006) and engage in crosstalk with myosin (Nishimura et al., 2021). In addition to elongation, formins appear to bundle F-actin, creating larger, thicker filaments (Esue et al., 2008). Both elongation and bundling allow FHOD1&3 and INF2 around the nucleus to generate an F-actin network that may protect the nucleus or at the very least coordinate nuclear movement (Antoku et al., 2023; Shao et al., 2015). Protection from mechanical insults is also mediated by DIAPH1 (Valencia et al., 2021). Finally, the formins FHOD3 and DIAPH3 appear to localize and modulate parallel and perpendicularly oriented stress fibers (Monzo et al., 2016; Tojkander et al., 2011). These distinct functions position formins as central players in cytoskeletal organization that leads to cell migration.

In particular, the formins FHOD3 and DIAPH3 have been implicated in cell morphology, cell migration and stress fiber formation. FHOD3 regulates F-actin is due to both its elongation and bundling activities (Antoku et al., 2019). FHOD3 overexpression facilitates migration and invasion in medulloblastoma through the RhoA/ROCK1/LIMK1 pathways (Yu et al., 2020). FHOD3 is phosphorylated by ROCK and overexpressed in ovarian cancer and its knock-down diminishes cell migration and invasion (Huang et al., 2025; Paul et al., 2015). This seems to place FHOD3 and ROCK in a positive feedback loop. This suggests a role of FHOD3 in regulating stress fibers. Indeed, FHOD3 is necessary for contact guidance, a directed migration strategy, through the production of parallel stress fibers (Monzo et al., 2016). It also controls 3D random migration through the formation of bundled F-actin spikes at the leading edge of cells that are parallel with the extended protrusion (Paul et al., 2015). DIAPH3 also controls cell morphology, cell migration and stress fiber formation. DIAPH3 loss drives cells from mesenchymal morphologies to amoeboid morphologies, down-regulating cell migration in cancer cells (Hager et al., 2012). However, DIAPH3 loss can also up-regulate cell migration as seen in cancer cells, where DIAPH3 seems to induce an epithelial to mesenchymal transition (Dong et al., 2018) or in epithelial cells that do not change their migration mode (Gupton et al., 2007). Lowering the DIAPH3 expression in epithelial cells results in slower migration due diminishing stable F-actin structures in the lamella (Gupton et al., 2007), which is supported by work showing DIAPH3 localizes to perpendicular F-actin fibers in the lamella (Tojkander et al., 2011). While FHOD3 and DIAPH3 have been shown to regulation morphology, migration and stress fiber formation, it is unclear if they shift the balancing between parallel and perpendicular stress fibers in cells.

In this work, we examine the expression level of formins across several different cell types with different morphological and migration characteristics. Analysis shows that while certain formins seem to correlate with and perhaps regulate cell spreading (DIAPH1 and INF2), a different set appears to regulate cell elongation (FHOD3 and DIAPH3). This differential control over cell elongation suggested that these formins might be modulators of the balance of perpendicular and parallel stress fibers. Knocking down FHOD3 and DIAPH3 levels using RNA interference indeed diminishes cell migration and elongation, by effecting the balance of stress fibers oriented perpendicular and parallel with cell alignment. FHOD3 primarily facilitates the formation of parallel stress fibers, while DIAPH3 primarily facilitates the regulation of perpendicular stress fibers within the cell. Our findings highlight how distinct formins control distinct F-actin networks to regulate cell morphology and migration.

## 2. Results

### 2.1 Extracellular matrix stiffness modulates cell spreading and elongation in different ways across different cells

To define how extracellular matrix (ECM) stiffness influences cell morphology, we fabricated collagen-coated polyacrylamide (PAA) hydrogels with defined elastic moduli (0.2, 2, and 20 kPa), alongside a glass substrate as a control. We analyzed a panel of cell lines representing diverse cell migration modes: mesenchymal (HFF and MDA-MB-231), epithelial (HaCaT), amoeboid (THP-1), and a migration mode-switching cell line (WM266.4). We imaged attached and spread cells away from the center of the coverslip. Qualitative analysis revealed striking cell type dependent differences in morphology in response to substrate stiffness (**Figure 1A**). For instance, MDA-MB-231 cells exhibited a highly elongated, spindle-like morphology with fine extensions on soft substrates (0.2 kPa PAA) but adopted a more rounded yet still spread morphology on stiff substrates (glass). In contrast, HFFs were rounded and less spread on soft substrates but became highly elongated and spread on stiff substrates, displaying prominent, well-aligned actin stress fibers. HaCaT, WM266.4, and THP-1 cells, however, showed minimal changes in overall morphology, remaining relatively less spread across all stiffness conditions (**Figure 1A** & **S1A**). We next quantified these observations by measuring cell area and aspect ratio. HFFs demonstrated a strong positive correlation between substrate stiffness and both spread area (**Figure 1B**) and aspect ratio (**Figure 1C**). MDA-MB-231 cells presented a more complex phenotype. Their spread area was modest on 0.2 through 20 kPa PAA substrates, but increased significantly on glass (**Figure 1D**). Their aspect ratio was most pronounced on soft substrates and decreased dramatically with increasing stiffness (**Figure 1E**). HaCaT cells showed a biphasic response in spread area with a maximum on 20 kPa (**Figure 1F**), similar to what was observed in WM266.4 cells (**Figure S1B**). Aspect ratio decreased with increasing stiffness, but only cells plated on glass showed statistically significant decrease compared to the soft substrate (0.2 kPa PAA) (**Figure 1G**). To synthesize these relationships, we plotted cell aspect ratio against cell area for different cells on various stiffness (**Figure 1H & S1F**). This analysis revealed three distinct phenotypic classes. Some cell lines were relatively stiffness-insensitive including HaCaT, WM266.4, and THP-1 cells. HFF cells showed positive correlation between aspect ratio and cell area and MDA-MB-231cells showed negative correlation between aspect ratio and cell area. Collectively, these results demonstrate that the morphological response to ECM stiffness is not a universal cell behavior but is fundamentally determined by cell-intrinsic mechanisms, leading to distinct stiffness sensing phenotypes.

**Figure 1:**
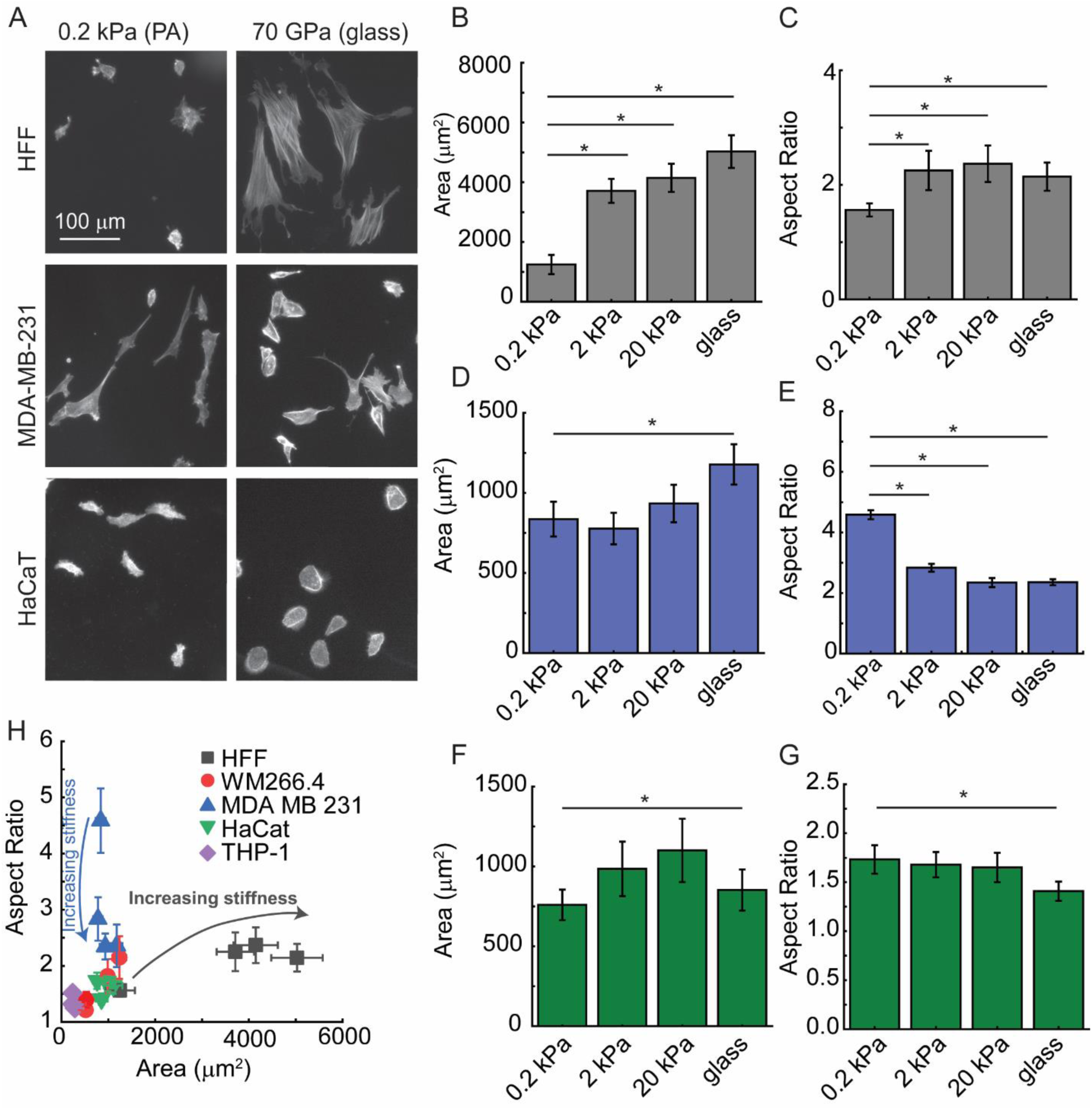
Matrix stiffness modulates cell spreading and elongation in different ways across different cells. (**A**) Representative fluorescence image of F-actin in HFF, MDA-MB-231 and HaCaT cells, illustrating distinct morphologies on 0.2 kPa PAA and glass substrates. (**B-G**) Quantification of (**B**, **D**, **F**) cell area and (**C**, **E**, **H**) cell aspect ratio for (**B**, **C**) HFF cells, (**D**, **E**) MDA-MB-231cells and (**F, G**) HaCaT cells on different substrate stiffnesses. (**H**) A scatter plot of cell aspect ratio versus cell area for all conditions reveals the stiffness-dependent morphological phenotype landscape. *N_samples_* = 3, *N_cells_* = 60 for each condition. Error bars represent a 95% confidence interval unless otherwise stated. *p*-values were calculated using a two-tailed unpaired Student’s *t*-test. * represents *p <* 0.005.

### 2.2 Extracellular matrix stiffness regulates cell migration across different cells

Having established that ECM stiffness dictates distinct morphological phenotypes in HFF, MDA-MB-231 and HaCaT cells (**Figure 1**), we next asked whether there were differences migratory behaviors. We hypothesized that the cell morphological response to the ECM would correspondingly define its migration response. We therefore quantified cell migration speed and persistence for these cell types plated on soft (0.2 kPa PAA) and stiff (glass) collagen coated substrates (**Figure 2A**). Representative migration trajectories revealed clear differences in migratory behavior (**Figure 2B-G**). HFF and HaCaT cells appeared slower on the stiff substrates, whereas MDA-MB-231 cells appeared persistent on the soft substrates. Quantitative analysis demonstrated that MDA-MB-231 cells underwent a shift in migration strategy on stiff substrates. Cell migration persistence time was significantly higher on soft substrates than on glass, while cell migration speed showed the opposite trend, significantly increasing on the rigid substrate (**Figure 2H&I**). In contrast, the migration of both HFF and HaCaT cells was largely unaffected by substrate stiffness, with no significant changes in either persistence time or speed between 0.2 kPa PAA and glass substrate (**Figure 2H&I**). To determine if a general relationship existed between these two migratory parameters, cell migration persistence time was plotted against speed (**Figure 2J**). We hypothesized that persistence time would correlate with aspect ratio and speed would anticorrelate with area (compare **Figure 2J** with **Figure 1H)**. However, while persistence time did seem to correlate with aspect ratio in MDA-MB-231 cells, stiff substrates that produced faster cells did not affect cell area. In addition, HFF cells did not show large changes in migration speed, even though stiffness changes resulted in large cell areas changes. This indicates that aspect ratio and persistence time are coupled, but not cell area and migration speed.

**Figure 2:**
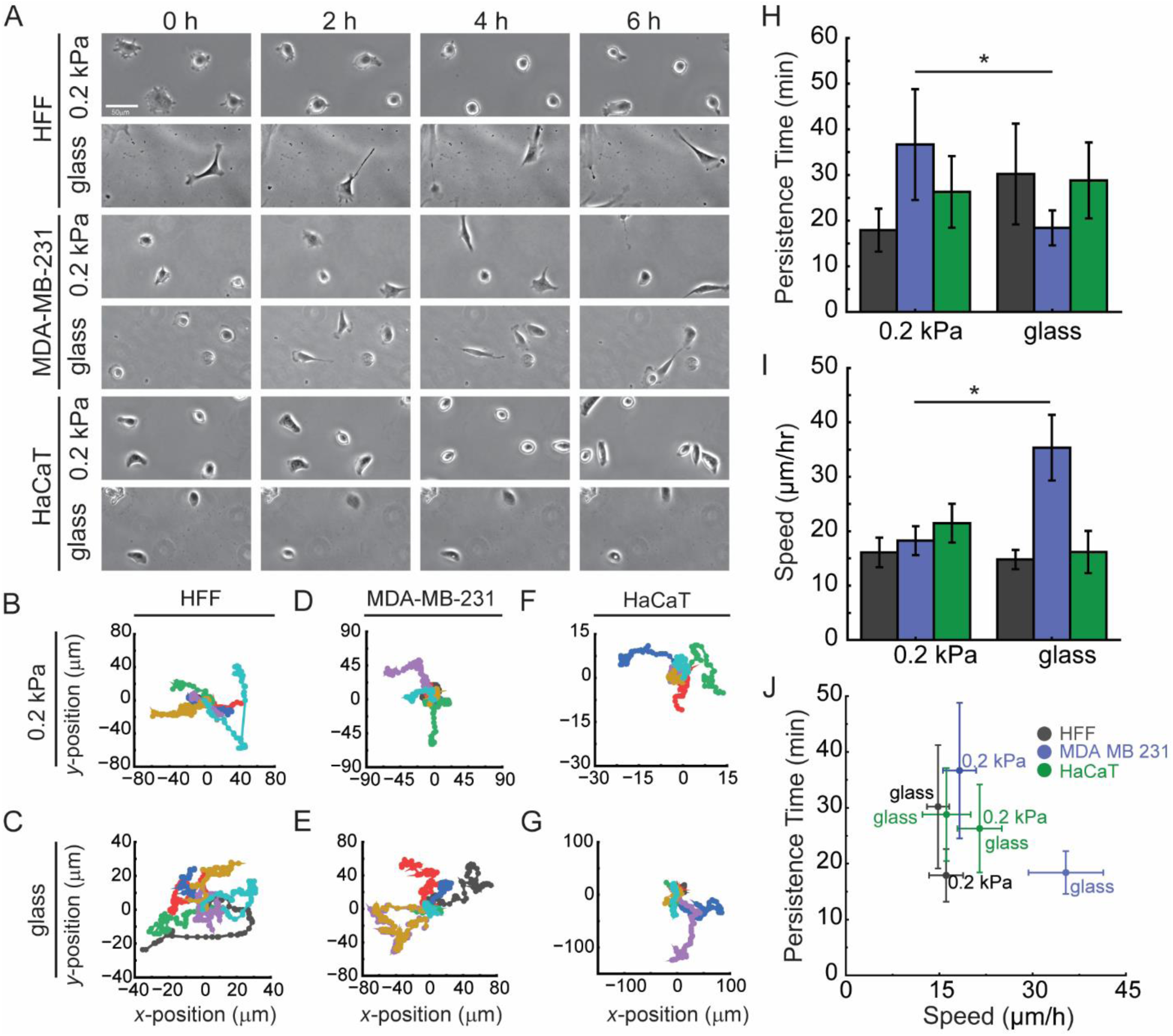
Extracellular matrix stiffness regulates cell migration across different cells. (**A**) Montages of live-cell phase contrast microscopy images (0, 2, 4 and 6 h) of HFF, MDA-MB-231, and HaCaT cells migrating on collagen-coated 0.2 kPa PAA and glass substrates. (**B-G**) Migration trajectories for individual cells with the initial position set at the origin and representing positions recorded every 5 min. (**B**) HFF on 0.2 kPa PAA, (**C**) HFF on glass, (D) MDA-MB-231 on 0.2 kPa polyacrylamide, (**E**) MDA-MB-231 on glass, (**F**) HaCaT on 0.2 kPa polyacrylamide, (**G**) HaCaT on glass (*N_tracks_* = 72). (**H, I**) Quantification of cell migration (**H**) persistence time and (**I**) speed as a function of substrate stiffness. (**J**) A scatter plot of cell migration persistence time versus cell migration speed for all conditions reveals the stiffness-dependent migration phenotype landscape (*N_samples_* > 4, *N_cells_*> 50). Error bars represent a 95% confidence interval unless otherwise stated. *p*-values were calculated using a two-tailed unpaired Student’s *t*-test. * represents *p <* 0.005.

### 2.3 Substrate stiffness governs F-actin cytoskeleton architecture across diverse cell morphologies and migration in different cells

Given the established role of linear F-actin structures in coordinating cell morphology and migration, we sought to determine whether the cell type specific differences we observed stem from distinct patterns of F-actin organization in response to stiffness. We hypothesized that substrate stiffness directs cell morphology and migration by differentially reorganizing the F-actin cytoskeleton across cell types. To test this, we performed a systematic quantitative analysis of stress fiber architecture in HFF, MDA-MB-231, and HaCaT cells on collagen-coated substrates of various stiffnesses. Fluorescence images of phalloidin-stained cells were processed using a local background subtraction method to isolate individual F-actin stress fibers for robust quantification (**Figure 3A&B**). We measured three key parameters: stress fiber density (number per cell), orientation angle (degrees) and length (μm). Stress fiber orientation was classified as parallel (aligned within 30° of the cell’s long axis), transition (30° - 60°), or perpendicular (60° - 90°) (**Figure 3C**). Strikingly, only HFFs assembled many prominent, long, and aligned actin stress fibers, and this was exclusively on stiff substrates (20 kPa PAA and glass). On soft (0.2 kPa) substrates, HFF F-actin networks were shorter and disorganized (**Figure 3D & S2B**). In contrast, both MDA-MB-231 and HaCaT cells failed to form robust, linear stress fibers regardless of substrate stiffness, maintaining only diffuse F-actin networks (**Figure 3D**). Quantification confirmed these observations. Stress fiber abundance was quantified by counting stress fibers longer than 5 μm. HFFs showed high and roughly similar stress fiber abundances, although there was a somewhat elevated level on 2 kPa PAA compared to other conditions. MDA-MB-231 and HaCat cells also exhibited a positive correlation between stress fiber abundance, albeit from a much lower baseline level of stress fibers (**Figure 3E**). The degree of orientation of stress fibers with respect to the long axis of the cell also depended on cell type and stiffness. HFF cells on glass exhibited a significant increase in stress fibers aligned with the cell direction (orientation angle < 30°) as stiffness increased, whereas MDA-MB-231 cells showed highest alignment on soft substrates (**Figure 3F**). No stress fiber orientation differences were observed on HaCaTs. Stress fibers were more aligned when aspect ratio was higher, irrespective of the stiffness that caused the high aspect ratio. Finally, stress fiber length was longest in HFF cells followed by MDA-MB-231 and HaCaTs and increased significantly with substrate stiffness in all three cell types (**Figure 3E**). In summary, the capacity to form a structured, aligned F-actin cytoskeleton in response to stiffness is a cell type-specific property. HFF cells, whose morphology and migration are strongly stiffness-dependent, assemble numerous highly ordered and long stress fibers on stiff substrates. MDA-MB-231 and HaCaT cells show increases in both stress fiber abundance and length with increased stiffness, but do not show more alignment with respect to the long axis of the cell.

**Figure 3:**
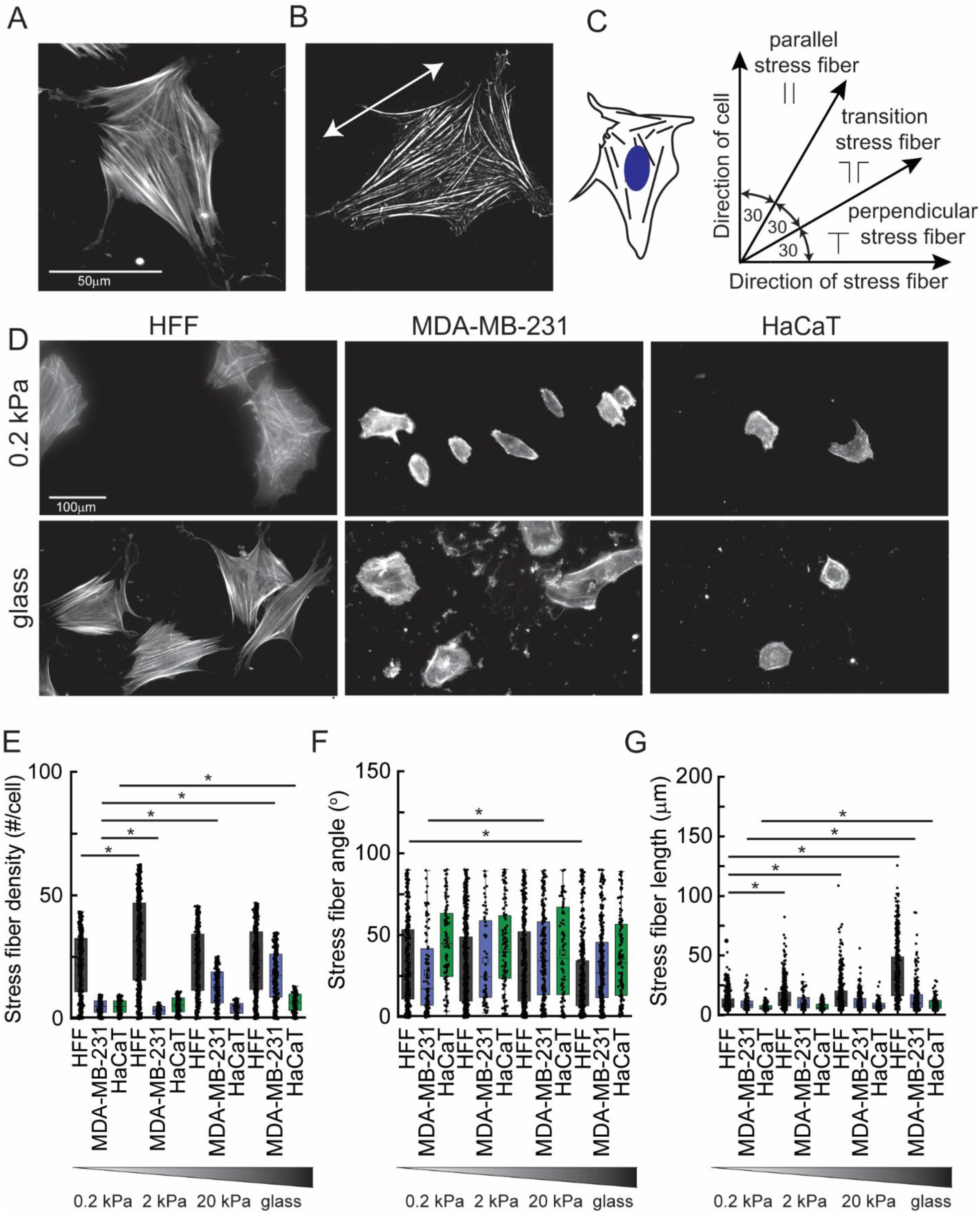
Substrate stiffness governs F-actin cytoskeleton architecture across diverse cell morphologies and migration modes. (**A**) A representative fluorescence image of F-actin in a spread cell shows stress fibers oriented in different directions. (**B**) A processed image after local background subtraction to resolve individual stress fibers for quantitative analysis. (**C**) Schematic defining stress fiber orientation angle relative to the cell’s major axis: parallel stress fibers (0° – 30°), transitional stress fibers (30° – 60°), perpendicular stress fibers (60° – 90°). (**D**) Representative fluorescence image of F-actin of HFF, MDA-MB-231, and HaCaT cells cultured on 0.2 kPa PAA and glass substrates. Quantification of cytoskeleton structure in response to stiffness: (**E**) Number of stress fibers per cell, (**F**) distribution of stress fiber orientation angle and (**G**) mean stress fiber length in different cell types on various substrate stiffness (*N_samples_* =3, *N_cells_* =10 and *N_SF_*> 66 and *N_cells_* =10). Box tops and bottoms refer to first and third quartiles (interquartile range. IQR), box midpoint lines refer to the median, and error bars (Whiskers) represent the minimum and maximum values within 1.5 times the IQR from the quartiles. All data are shown. *p*-values were calculated using a two-tailed unpaired Student’s *t*-test. * represents *p <* 0.005.

### 2.4 Different sets of formins correlate with cell area and aspect ratio

Cell type-specific differences in F-actin organization prompted us to investigate the molecular regulators that could underline this phenotypic variation across cell lines. Given their central role in nucleating and elongating linear F-actin fibers, we hypothesized that the expression of formins dictates the capacity for cells to generate stress fibers. We measured the transcript levels of all fifteen formin isoforms across our panel of cell types. Agarose gels show the correct RT-PCR product size and reveal differences in expression level across cells (**Figure S3**). For a more quantitative analysis, RT-qPCR was conducted and expression was normalized to GAPDH expression levels. Formins measured by RT-PCR and RT-qPCR showed similar expression level variability across cell lines (**Figure 4A**). Since we observed differences in stress fiber structure, cell migration and cell shape across cells, we hypothesized that formins which are both highly expressed as well as variable across the cell lines are good candidates for the controlling stress fiber structure, cell migration and cell shape. To identify formins that are both highly expressed as well as variable in their expression across the cell lines, we plotted the standard deviation of formin expression level against the average formin expression level (**Figure 4B**). The following formins were identified as strong candidates: DIAPH1, DIAPH2, DIAPH3, FHOD3, DAAM1 and INF2. However, in order to identify formins that specifically regulate different morphological features, we used Partial Least Squares (PLS) regression coupled with Variable Importance in Projection (VIP) scoring. This multivariate analysis identified formins whose expression levels are more strongly associated with variations in cell area (**Figure 4C**) and aspect ratio (**Figure 4D**) across all stiffness conditions. Stronger than average correlations occur when VIP > 1 and are denoted by the light grey shading. Candidates identified in **Figure 4B** are shown in dark grey shading. The PLS VIP analysis revealed a striking segregation of function. Cell spread area was most strongly associated with DIAPH1, DAAM2, FMN2, FMNL2, INF1, and INF2 (**Figure 4C**). In contrast, cell aspect ratio was most strongly associated with DIAPH1, DIAPH3 and FHOD3 (**Figure 4D**). Only DIAPH1 seemed to be linked to both cell area and aspect ratio. These findings suggest that different sets of formin family members regulate distinct aspects of cell morphology, potentially through the assembly of specialized F-actin networks.

**Figure 4:**
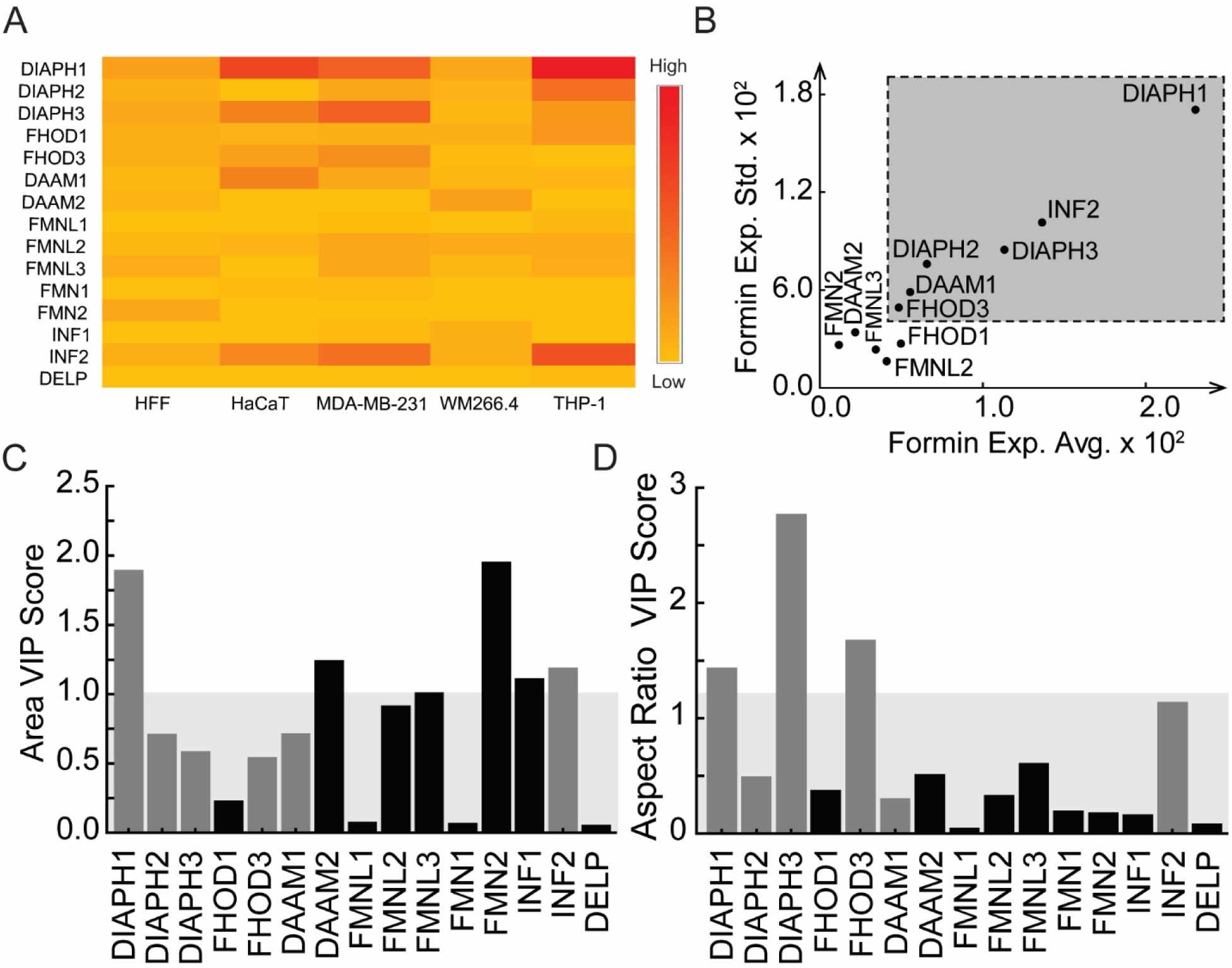
Different sets of formins correlate with cell area and aspect ratio. (**A**) Heatmap of formin expression normalized to GAPDH across five human cell types. Formins are grouped based on their sequence homology. (**B**) The standard deviation of formin expression level plotted against the mean formin expression level normalized to GAPDH across five cell lines. The grey box indicates formins that are both highly expressed and variably expressed across the five cell lines. Variable importance in projection (VIP) scores from a partial least squares (PLS) regression model linking formins expression to (**C**) cell area and (**D**) cell aspect ratio. VIP scores over 1 (light grey region) indicate the particular formin is more important than the average formin in predicting the variation in the morphological parameter. Formins that are found in the upper right area of the graph in (**B**) are shown in grey. Those with low levels of expression and/or low variability across cells are shown in black. *N_biological replicates_* = 2, *N_technical replicates_* = 2 and *N_total_*= 4.

### 2.5 FHOD3 and DIAPH3 regulate cell morphological properties in different cells

The PLS VIP analysis identified FHOD3 and DIAPH3 as formin family members that most strongly associate with cell aspect ratio (**Figure 4D**). To determine if this correlation reflects a functional requirement, we investigated whether these formins directly regulate the morphological features of HFFs and MDA-MB-231 cells on soft (0.2 kPa) and stiff (glass) substrates. We performed siRNA-mediated knockdown (KD) of FHOD3 and DIAPH3 in both cell types (**Figure 5A**). Transcript level was diminished in HFF cells 4-5 fold (**Figure 5B&C**) and in MDA-MB-231 cells 2-fold (**Figure 5D&E**). The KD of FHOD3 resulted in shorter extensions and accumulation of an F-actin rich periphery in MDA-MB-231 cells, whereas HFF cells were larger on soft substrates and smaller on stiff substrates (**Figure 5A**). The KD of DIAPH3 showed similar responses, although there was a less-pronounced F-actin rich periphery in MDA-MB-231 cells. Cell area and aspect ratio were quantified (**Figure 5F&G**). In HFF cells on soft substrates, FHOD3 KD increased cell area while DIAPH3 KD decreased it. On glass substrates, however, both KD conditions led to a significant reduction in HFF spread area (**Figure 5F**). In MDA-MB-231 cells on soft substrates, both FHOD3 and DIAPH3 KD result in smaller cell area and on stiff substrates, no change was observed (**Figure 5F**). Changes in aspect ratio were the same for soft substrates. In HFF cells, FHOD3 KD increased cell aspect ratio while DIAPH3 KD decreased it, while in MDA-MB-231 cells, both FHOD3 and DIAPH3 KD diminished aspect ratio (**Figure 5G**). On stiff substrates the cell type behavior was switched. In HFF cells, FHOD3 and DIAPH3 KD did nothing, whereas in MDA-MB-231 cells, both FHOD3 and DIAPH3 KD diminished aspect ratio (**Figure 5G**). These data demonstrate that FHOD3 and DIAPH3 do not simply govern cell elongation as was hypothesized, but rather alter both cell area and aspect ratio. While FHOD3 and DIAPH3 KD result in similar decreases in both cell area and aspect ratio, the response of HFFs on soft substrates seem to indicate differential control by FHOD3 and DIAPH3 over cell morphology.

**Figure 5:**
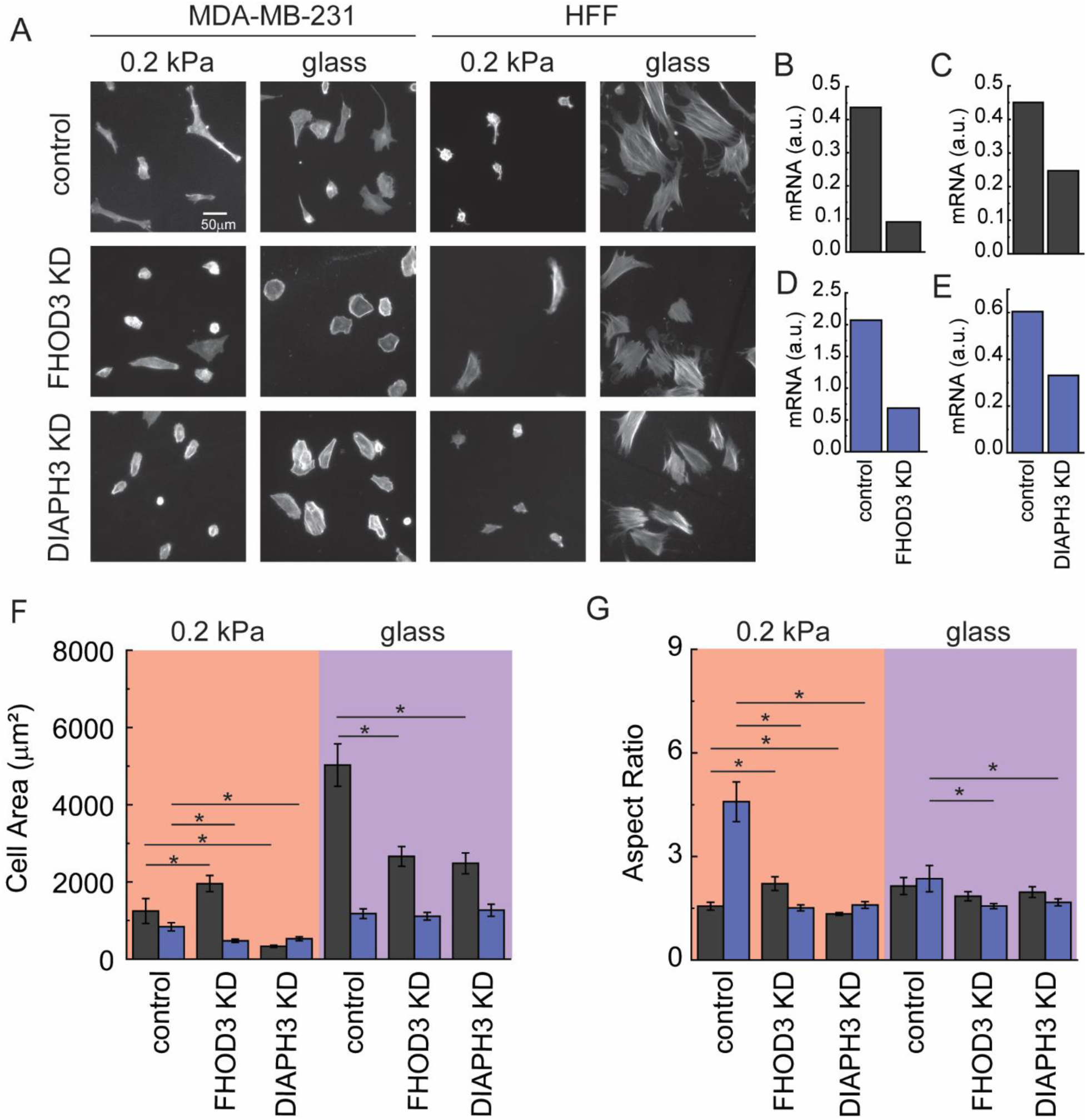
FHOD3 and DIAPH3 regulate cell morphological properties in different cells. (A) Representative immunofluorescence image of F-actin(phalloidin-568) in control, FHOD3KD, and DIAPH3KD in MDA-MB-231 and HFF cells plated on 0.2 kPa PAA and glass substrates. Scale bar is 50μm. (B, C), and (D,E), Reverse Transcription Polymerase chain reaction analysis of *FHOD3*(B) and Quantitative RT-PCR analysis of DIAPH3 mRNA expression in HFF and MDA-MB-231 cells following KD, normalized to GAPDH.(F,G) Measurement of cell aspect ratio (F) and area(G) in control, FHOD3KD, and DIAPH3KD in cells cultured on collagen coated on 0.2 kPa PAA and glass substrates. *N_cells, HFF_* > 50, and *N_cells, MDA-MB-231_* >50. Error bars represent a 95% confidence interval unless otherwise stated. P-values were calculated using a two-tailed unpaired Student’s t-test. (Significant * represents *p <* 0.005. All the experiments were replicated at least three times unless otherwise stated.

### 2.6 FHOD3 and DIAPH3 govern cell migration in different cells

Having established that FHOD3 and DIAPH3 are critical regulators of cell morphology, we next asked whether they similarly govern cell migration. We quantified the migration speed and persistence of HFFs and MDA-MB-231s following siRNA-mediated KD of FHOD3 and DIAPH3 on soft (0.2 kPa) and stiff (glass) substrates (**Figure 6A and S4A**). The impact of formin depletion was cell type-specific. In HFFs, KD of neither FHOD3 nor DIAPH3 significantly altered cell migration speed or persistence time on either substrate (**Figure 6B&C**). In contrast, MDA-MB-231 cells exhibited a strong dependence on both formins for motility on both substrates. KD of either FHOD3 or DIAPH3 significantly reduced cell migration speed on both soft and stiff substrates (**Figure 6B&D** and **S4B**). Despite this reduction in speed, no statistically significant change in migration persistence in both HFF and MDA-MB-231 cells with either formin KD was observed (**Figure 6C&D** and **S4B**). This was somewhat surprising given that both formins induced robust decreases in aspect ratio (**Figure 5G**). These functional studies demonstrate that FHOD3 and DIAPH3 are required for high speeds, but only in highly motile, MDA-MB-231 cells. In less motile cells, FHOD3 and DIAPH3 affect morphology and stress fiber structure, but not migration speed.

**Figure 6.**
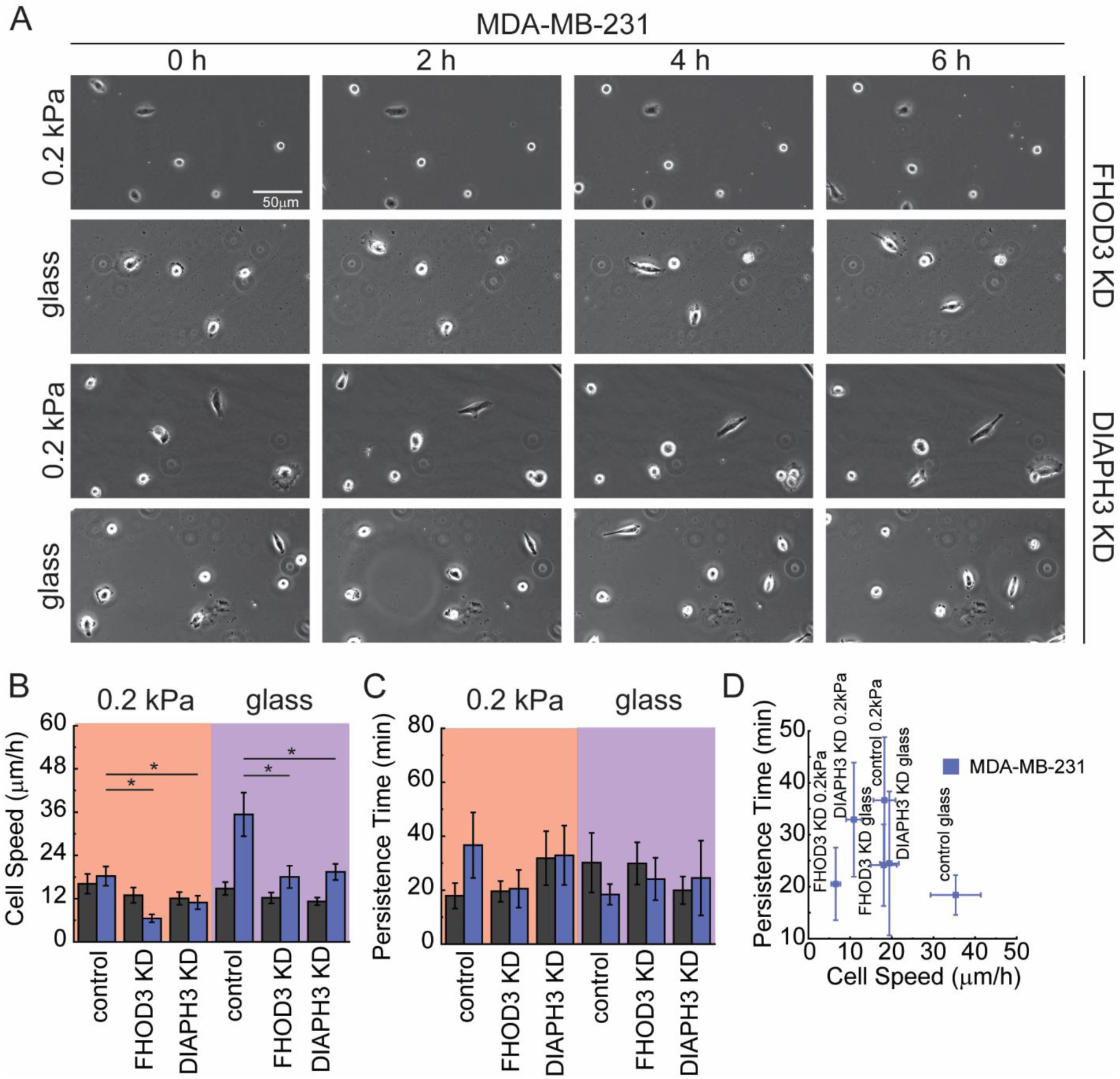
FHOD3 and DIAPH3 govern cell migration in different cells. (A) Representative phase contrast time-lapse montage of MDA-MB-231 cells with FHOD3KD and DIAPH3KD on 0.2 kPa PAA and glass substrates. Scale bar:50μm. (B, C) Quantifying cell migration speed (B) and (C) persistence time for control, FHOD3KD, and DIAPH3KD in HFF and MDA-MB-231 cells. (D) Scatter plot illustrating the relationship between cell speed and persistence time for all conditions and substrates. *N_cells, HFF_* > 55, and *N_cells, MDA-MB-231_* >55. Error bars represent a 95% confidence interval unless otherwise stated. P-values were calculated using a two-tailed unpaired Student’s t-test. (Significant * represents *p <* 0.005. All the experiments were replicated at least three times unless otherwise stated.

### 2.7 FHOD3 and DIAPH3 shift the balance between parallel and perpendicular stress fibers

Having established that FHOD3 and DIAPH3 regulate cell morphology and migration (**Figure 5&6**), we sought to quantify the effect on different stress fiber populations. When cells exhibit an elongated morphology, they produce many stress fibers that are parallel with the elongation axis, but also those that are perpendicular to the elongation axis. The relative numbers and lengths likely define the morphology. We hypothesized that FHOD3 and DIAPH3 orchestrate the organization of different stress fiber populations. To test this, we imaged stress fiber architecture in HFF and MDA-MB-231 cells following FHOD3 or DIAPH3 KD on soft (0.2 kPa) and stiff (glass) substrates (**Figure 7A**). We first quantified stress fiber length. Knock-down of either FHOD3 or DIAPH3 resulted in lower average stress fiber length, supporting the role of these fomins in linear F-actin elongation (**Figure 7B**). In addition to average stress fiber length, we computed a total stress fiber length, which is the sum of the lengths of all the stress fibers and constitutes and amount of actin incorporated into stress fibers (**Figure 7C**). Again, there were large decreases in the amount of actin incorporated into stress fibers when either FHOD3 or DIAPH3 were knocked down. We examined the average orientation angle of the fibers with respect to the long axis of the cell, where an angle of 0 refers to stress fibers that are well aligned in the direction of cell alignment and higher angles refer to more disorganized alignment (**Figure 7D**). Unlike stress fiber length, stress fiber orientation showed formin-specific differences. FHOD3 KD in HFF cells on glass showed poorer stress fiber orientation, whereas DIAPH3 KD showed better stress fiber orientation. MDA-MB-231 cells showed a different response with DIAPH3 KD, as it increased stress fiber orientation. Changes in average orientation suggest changes in parallel and perpendicular stress fiber populations, so we assessed total stress fiber length in either parallel or perpendicular orientation in response to FHOD3 and DIAPH3 KD in MDA-MB-231 and HFF cells (**Figure 7E&F**). In MDA-MB-231 cells both formins caused a decrease in total length, but the parallel stress fibers seemed to be more sensitive (**Figure 7E**). In HFF cells both formins caused a decrease in total length of parallel stress fibers. However, on glass FHOD3 caused an increase in total length of perpendicular stress fibers, whereas DIAPH3 caused a decrease in total length of perpendicular stress fibers. To analyze the change in the balance of total stress fiber length in the parallel and perpendicular direction, the ratio of total stress fiber length was calculated (**Figure 7G**). In MDA-MB-231 cells on soft substrates, KD of either formin diminished the parallel to perpendicular stress fiber balance, whereas on glass there was no change. In HFFs cells, no change was observed on soft substrates, but a large swing was present on glass. FHOD3 KD dramatically diminished the parallel to perpendicular ratio, whereas DIAPH3 KD dramatically increased the parallel to perpendicular ratio (**Figure 7G**). In summary, our data suggests that FHOD3 and DIAPH3 regulate linear F-actin stress fibers through complementary yet distinct mechanisms. Both act to elongate stress fibers, but FHOD3 acts to foster parallel stress fibers, whereas DIAPH3 acts to foster perpendicular stress fibers. This indicates that formin family members tune the balance of different populations of stress fibers with cells.

**Figure 7:**
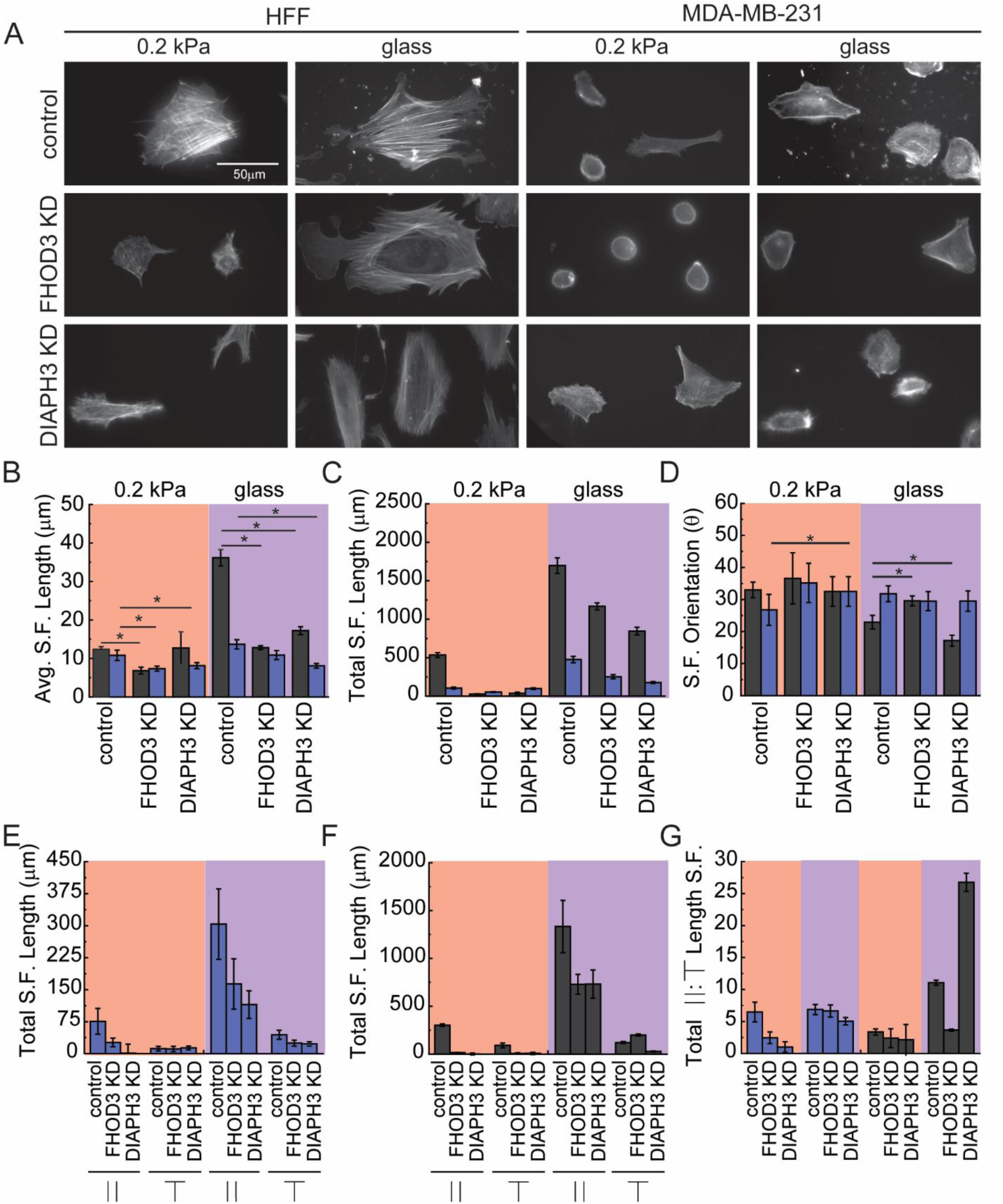
FHOD3 and DIAPH3 govern stress fiber structure through complementary, yet distinct molecular mechanisms. (A) Representative immunofluorescence images of F-actin (phalloidin-568) in control, FHOD3KD, and DIAPH3KD in HFF and MDA-MB-231 cells plated on 0.2kPa PAA and glass substrates. Scale bar:100μm. (B, C) Quantification of mean stress fiber length (B) and total stress length (C) for the indicated conditions. (D) Distribution of stress fiber angles(θ) relative to the cell’s major axis, categorized as parallel stress fibers (0°–30°) or perpendicular (60°–90°). (E, F) Quantifying total parallel and perpendicular stress fiber length in MDA-MB-231 cells (E) and HFF cells (F) in control, FHOD3KD, and DIAPH3KD on 0.2kPa PAA and glass. (G) Ratio of parallel to perpendicular stress fibers in MDA-MB-231 and HFF cells across substrates and conditions on 0.2kPa PAA and glass substrates. *N_cells, HFF_* =10, and *N_cells, MDA-MB-231_* >10, 25 < *N_Stress Fibers_* < 201, Error bars represent a 95% confidence interval unless otherwise stated. P-values were calculated using a two-tailed unpaired Student’s t-test. (Significant *represents *p <* 0.005. All the experiments were replicated at least three times unless otherwise stated.

**Figure 8:**
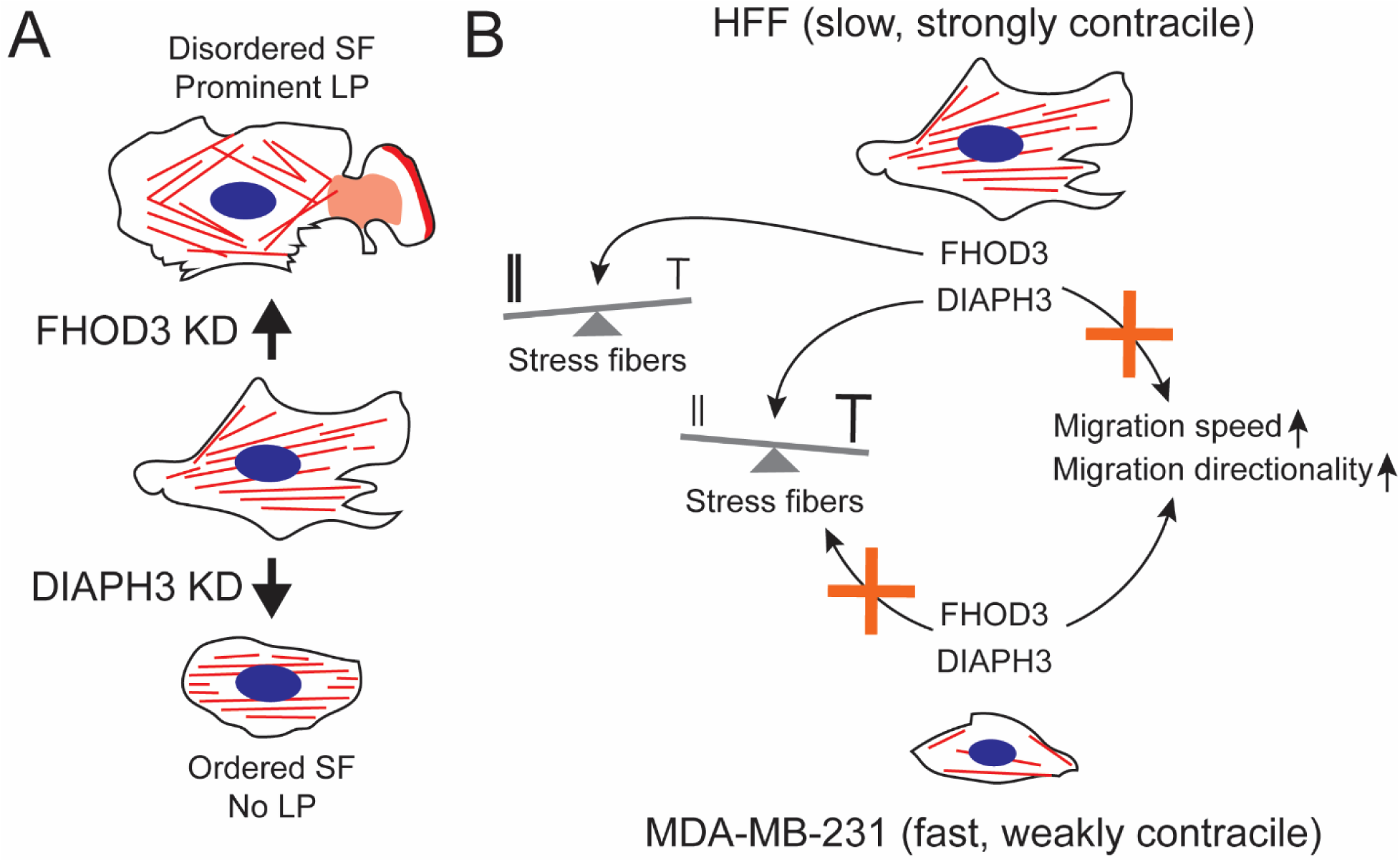
Schematic outlining the role of FHOD3 and DIAPH3 in governing F-actin structure and cell migration. (A) Role of FHOD3 and DIAPH3 in altering F-actin structure at the level of stress fibers (SF) and lamellipodium (LP). (B) Differing roles of FHOD3 and DIAPH3 in controlling F-actin structure and cell migration in HFFs and MDA-MB-231s.

## 3. Discussion

In this paper we have explored the role of different formins in regulating cell shape and cell migration through the assembly of different populations of F-actin stress fibers. We use extracellular matrix (ECM) stiffness as a way to promote a diversity of cell shapes across different cell types. Cells can adopt a variety of morphologies that are dependent on stiffness. However, stiff ECM does not always result in similar changes in cell elongation and cell area across all cell types. Stiff environments can make cells shorter (MDA-MB-231), while keeping the area roughly the same or make cells larger (HFFs), while keeping the aspect ratio roughly the same. Alternatively, stiff environments can make cells both longer and larger (WM266.4 and HaCat). The spreading of cells on the softest substrates seemed somewhat surprising, given literature indicating that cells do not spread well on soft substrates. However, we chose to image cells close to the edge of the substrate, where the gel might be stiffer, due to our desire to analyze spread and elongated cells. We also measured cell migration in these regions around the edge. We hypothesized that persistence time would be proportional to aspect ratio and it is across several cell types. We also hypothesized that cell migration speed would be inversely proportional to cell area. It was not. Smaller cells could be either faster or slower than more spread cells. In our hands the fastest cells were the relatively elongated MDA-MB-231 cells (**Figure 2I**). Therefore, we focused on formins that regulated elongation most strongly (**Figure 4D**). The expression of the formin genes FHOD3 and DIAPH3 (whose protein product is confusingly called mDia2 and Dia2) showed the largest correlation with aspect ratio across the cell types. Increased expression of FHOD3 upregulates migration in glioblastoma cells (Heuser et al., 2020). Additionally, FHOD3 upregulates integrin-mediated cell migration in 3D collagen matrices (Paul et al., 2015) as well as contact guidance, directed migration on lines of ECM (Monzo et al., 2016). Mesenchymal mode cell migration is highly integrin dependent and mesenchymal cells robustly contact guide. Our results showing FHOD3 knockdown (KD) decreases cell migration speed in mesenchymal cells across different stiffnesses agrees with this (**Figure 6B**). DIAPH3 is also a key regulator of cell migration through its role in assembling contractile actin structures in the lamella, particularly transverse arcs (Gupton et al., 2007; Tojkander et al., 2011). These fibers are essential for generating the actomyosin contractility that powers lamellar contraction resulting in regulation of protrusion, focal adhesion strengthening and body translocation (Gupton et al., 2007). DIAPH3 has mixed migration effects. It tends to induce a mesenchymal mode of cell migration, which can either increase migration in epithelial cells (Dong et al., 2018; Gupton et al., 2007) or decrease migration in amoeboid cells (Hager et al., 2012). Our results show DIAPH3 KD decreases cell migration speed in MDA-MB-231 cells (**Figure 6B**), which agrees with previous work suggesting it is a positive regulator of migration (Dong et al., 2018; Gupton et al., 2007). However, DIAPH3 KD also seems to reduce spreading of MDA-MB-231 cells, perhaps suggesting that DIAPH3 is important in retaining the mesenchymal mode of migration. However, we don’t see increases in cell migration as was observed elsewhere (Hager et al., 2012). Consequently, we observe robust down-regulation of migration when both FHOD3 and DIAPH3 are knocked down.

As mentioned previously, we focused on understanding how formins regulate elongation as measured by aspect ratio. Spread, elongated mesenchymal cells (MDA-MB-231) tend to migrate faster than cells that are spread, but lack elongation (HFF, **Figure 2I**). Furthermore, elongation may allow mesenchymal cells in certain circumstances such as cancer metastasis to move through tight spaces (Paul et al., 2017). Finally, directional cues like contact guidance result in elongated mesenchymal cells that can both migrate with higher directional persistence and remodel collagen fibers to produce alignment (Barocas and Tranquillo, 1997; Wang and Schneider, 2017). To identify key regulators of cell elongation, we employed a Variable Importance in Projection (VIP) analysis, which quantifies the correlation between formin expression patterns and morphological features across cell lines. The two formins with the largest correlation with aspect area were FHOD3 and DIAPH3. Both have roles in elongation. FHOD3 has been a demonstrated regulator of integrin-dependent protrusion (Paul et al., 2015) and contact guidance (Monzo et al., 2016), which suggests its role in promoting an elongated phenotype. DIAPH3 expression seems to promote the mesenchymal phenotype typified by extensions from either amoeboid cells (Hager et al., 2012) or epithelial cells (Dong et al., 2018). DIAPH3 also has been associated with stabilizing lamellipodial extension (Gupton et al., 2007; Yang et al., 2007). While we observe that the KD of FHOD3 and DIAPH3 produce less elongated cells (**Figure 5G**), this is only true across both soft and stiff ECM in MDA-MB-231 cells. In HFFs, FHOD3 and DIAPH3 KD did little to cell elongation on stiff substrates. It appears to diminish elongation slightly, but this was not statistically significant. However, HFFs have relatively small aspect ratios so the assay may not have the ability to detect changes in an already small number (**Figure 5G**). On soft substrates, FHOD3 KD actually produced more elongated cells (**Figure 5G**). It is possible that KD of FHOD3 limits Rho activity and diminishes myosin contractility (Yu et al., 2020), which in control HFFs may normally cause deadhesion on soft ECM due to an improper balance between contractility and adhesion. Indeed, cell area also increases in response to FHOD3 KD (**Figure 5F**). DIAPH3 has no such effect on Rho activity and thus its KD causes decreases in both cell area and aspect ratio. Taken together this shows an important role for FHOD3 and DIAPH3 in regulating cell elongation.

Given that FHOD3 and DIAPH3 seem to regulate elongation, we were interested if they also modulated F-actin stress fiber formation. Recall that FHOD3 promotes F-actin polymerization and bundling (Antoku et al., 2019) and localizes to parallel F-actin stress fibers, likely ventral stress fibers (Monzo et al., 2016). These are connected to F-actin spikes that are required for integrin-mediated cell migration and are largely oriented parallel with cell migration (Paul et al., 2015). FHOD3 activates and is activated by ROCK in a positive feedback loop (Paul et al., 2015; Yu et al., 2020). The activation of ROCK by FHOD3 suggests that it encourages the assembly of stress fibers through myosin activation as well as diminishes lamellipodial activity, potentially by diminishing branched F-actin network through the inhibition of cofilin by LIMK (Mouneimne et al., 2006). This explains both the lack of ventral stress fibers as well as the enhanced presence of lamellipodia in HFFs (**Figure 7A**). DIAPH3 on the other hand seems to promote F-actin polymerization in the lamella (Gupton et al., 2007) and localizes to perpendicular F-actin stress fibers that are likely transverse arcs (Tojkander et al., 2011). In HFFs on stiff substrates stress fiber lengths go down (**Figure 7B**) and stress fiber numbers go up (**Supplementary Figure 7D**) with either FHOD3 or DIAPH3 KD, suggesting that elongation and not nucleation is the driving function of both of these formins on stiff environments. MDA-MB-231 have fewer and shorter stress fibers and FHOD3 and DIAPH3 KD seem to diminish both stress fiber length and numbers suggesting regulation over both elongation and nucleation. What is most interesting are results associated with the control by formins over parallel and perpendicular stress fibers. We characterized stress fiber polymerization with total stress fiber length, which accounts for both nucleation and elongation. Both FHOD3 and DIAPH3 KD diminish parallel stress fiber polymerization in HFFs and MDA-MB-231s. However, in HFF cells FHOD3 KD results in more perpendicular stress fiber polymerization and DIAPH3 results in less perpendicular stress fiber polymerization. The consequence is that for HFFs with large stress fibers FHOD3 KD tips the scales towards perpendicular stress fiber polymerization, whereas DIAPH3 KD tips the scales towards parallel stress fiber polymerization (**Figure 7G**). This is not seen in MDA-MB-231 where the ratio of parallel to perpendicular is unaffected by FHOD3 or DIAPH3 KD. It is not known whether this change in the balance of parallel and perpendicular stress fibers is caused by the action of other F-actin regulatory proteins that are not present in HFFs or is due to the fact that MDA-MB-231s are less contractile than HFFs (Foroozandehfar et al., 2025) and thus have a much lower number of detectable stress fibers. While the HFF response of the ratio of parallel to perpendicular stress fibers maps well with previous work that links FHOD3 to parallel (Monzo et al., 2016) and DIAPH3 to perpendicular (Tojkander et al., 2011) stress fibers, there seems to be a more complicated story. It is possible that free G-actin or some other F-actin regulatory protein that is not used for parallel stress fiber polymerization under FHOD3 KD is used for DIAPH3-mediated perpendicular stress fibers, resulting in the increase in perpendicular stress fiber formation. DIAPH3 could potentially regulate parallel stress fibers that reside in the lamella, a region where DIAPH3 is working. Consequently, FHOD3 and DIAPH3 have opposing effects on the balance of parallel and perpendicular stress fibers in highly contractile cells (HFFs), but do not alter the balance of stress fibers in highly motile cells (MDA-MB-231).

The fact that FHOD3 and DIAPH3 rebalance parallel and perpendicular stress fibers might have an interesting impact on how cells transmit forces to the ECM (Niraula et al., 2025) or exert internal tension (Amiri et al., 2023). We predict that FHOD3 and DIAPH3 cause a rebalancing of traction stresses in the parallel and perpendicular directions that likely affects how cells remodel ECM or spread and migrate, particularly on aligned collagen fibers. FHOD3 promotes the formation of parallel, ventral stress fibers (Monzo et al., 2016), which are primary generators of high-magnitude axial traction forces and are associated with contact guidance along the length of aligned collagen fibers. These parallel, ventral stress fibers also seem to bear large internal cytoskeletal tension (Lee et al., 2018), particularly when cells are extended (Amiri et al., 2023). In contrast, DIAPH3 favors the assembly of perpendicular stress fibers or transverse arcs (Tojkander et al., 2011) that might have the effect of radially pulling sides of the cell inward, controlling cell shape by making the cell thinner. This thinning of the cell along with axial elongation would create a high aspect ratio shape that is primed for directed migration along aligned ECM. This radial force generation inward could also act to pull collagen fibrils closer and help the cell to bundle them into collagen fibers. We propose that functional specialization allows these formins to differentially govern mechanical output to ECM. Consistent with this model, perturbations in HFFs confirm their opposing roles. FHOD3 KD shifts the cytoskeletal balance toward perpendicular stress fibers and likely reduces axial contractility, whereas DIAPH3 KD enhances parallel fiber dominance and likely increases radial tension (**Figure 7G**). Consequently, FHOD3 depletion which diminishes parallel fibers is predicted to reduce traction forces in the direction of collagen alignment, while potentially elevating axial traction, even though it has little effect on the average traction stress magnitude (Niraula et al., 2025). Conversely, we predict that DIAPH3 KD, by enhancing parallel stress fiber polymerization, likely increases traction anisotropy and results in less radial traction stress and perhaps more axial traction stress. Thus, in regulating the balance of parallel and perpendicular stress fibers, FHOD3 and DIAPH3 may be able to balance traction stresses exerted in different directions. This would allow the cell to do two things. First, this balancing mechanism would allow for control over the generation of forces that reinforce existing collagen alignment via parallel stress fibers or promote fiber realignment via perpendicular stress fibers, thus regulating ECM remodeling. Second, this mechanism would allow the cell to control forces that both regulate cell shape and pull the cell forward, particularly in response to aligned ECM during contact guidance. In conclusion, FHOD3 and DIAPH3 could regulate traction forces exerted in different directions by facilitating the assembly of different populations of stress fibers.

## 4. Conclusion

In this study, we provide a systematic analysis of how the formin family of actin regulatory proteins governs cell morphology, migration and F-actin cytoskeletal structure. We first establish that cell types with different migration modes including mesenchymal, epithelial and amoeboid exhibit distinct morphological and cytoskeletal responses to substrate stiffness, where elongated cells migrated most quickly. We measured formin expression across these cell types and employed multivariate regression, to identify FHOD3 and DIAPH3 potential regulators of cell elongation. KD studies revealed that these two formins regulate cell morphology, migration, and stress fiber populations. In highly migratory cells (MDA-MB-231s), both FHOD3 and DIAPH3 KD diminished cell migration, but had little impact on the F-actin stress fiber alignment and the balance between parallel and perpendicular stress fiber polymerization. In less migratory cells (HFFs), both FHOD3 and DIAPH3 knock-down did not affect cell migration, but had a dramatic impact on the F-actin stress fiber alignment and the balance between parallel and perpendicular stress fiber polymerization. FHOD3 KD acted to diminish stress fiber alignment with the long axis of the cell by both diminishing parallel stress fiber polymerization and increasing perpendicular stress fiber polymerization. DIAPH3 KD acted to increase stress fiber alignment with the long axis of the cell by dramatically diminishing perpendicular stress fiber polymerization, while only diminishing parallel stress fiber polymerization. The results indicate that FHOD3 and DIAPH3 control the balance over parallel and perpendicular stress fiber polymerization, allowing the cell to tune the cytoskeletal structure for the control over cell morphology and migration.

## 5. Materials and Methods

### 5.1 Cell lines and culturing

A human mammary carcinoma cell line (MDA-MB-231, ATCC, Manassas, VA), an immortalized human foreskin fibroblast cell line (HFF, ATCC), an immortalized human keratinocyte cell line (HaCaT, kind gift from Dr. Torsten Wittmann), a metastatic human melanoma cell line (WM-266-4, ATCC) and an immortalized monocytic cell line (THP-1, ATCC) were used in this study. MDA-MB-231, HaCaT and WMM266-4 cells were cultured in Dulbecco’s Modified Eagle’s Medium (Sigma Aldrich, St. Louis, MO), supplemented with 10% fetal bovine serum (Gibco). HFFs were supplemented with 15% fetal bovine serum (Gibco), 1% penicillin-streptomycin (Gibco), and 1% Glutamax (Gibco) and THP-1 cells were cultured in RPMI-medium 1640 1x (Gibco) with 10% FBS, 1% penicillin streptomycin, 0.05 mM BME (β-Mercaptoethanol, Cas No: 60-24-2, (Sigma Aldrich). at 37 °C in 5% CO_2_. When live cells were imaged, cells were placed in culture media that lacked phenol red, but contained 15 mM 4-(2-hydroxyethyl)-1-piperazineethanesulfonic acid, (Sigma Aldrich,), 1% Glutamax, 1% penicillin/streptomycin and 10% or 15% FBS, depending on the cell line. This is referred to as clear imaging media.

### 5.2 Fabrication of polyacrylamide gels and glass substrates

This study used polyacrylamide (PAA) gels to generate collagen substrates of different stiffness. First, 22 x 22 mm coverslips were thoroughly cleaned and then covered with 0.1 M NaOH for 10 min. The coverslips were then treated with a 5% 3-aminopropyltrimethoxysilane (APTMS, Thermo Scientific, Waltham, MA) solution for 5 min. The coverslips were then washed three times with deionized water, each lasting 5 min. Next, a 0.5% glutaraldehyde solution (Electron Microscopy Sciences, Hatfield, PA) was applied to the coverslips for 30 min, followed by additional washes with deionized water. To prepare PAA gels with different elastic moduli, solutions of 3%, 5% or 8% acrylamide (Bio-Rad, Hercules, CA) were added to solutions of 0.03%, 0.1%, or 0.26% bis-acrylamide (Bio-Rad), respectively. In addition, 0.05% Ammonium Persulfate (APS, Bio-Rad) and 0.15% N,N,N’, N’-Tetramethylethylenediamine (TEMED, Fisher Scientific), respectively, to achieve gels with elastic moduli of 0.2, 2, and 20 kPa. The functionalized coverslips were placed face-down onto drops of these solutions on microscope slides, allowing polymerization to occur for 20 min. The samples were then treated with 2 mM Sulpho-SANPAH (Thermo Fisher) under UV light using a 365 nm light source at a power of 3 mW for 8 min to functionalize their surfaces. The PAA substrates on the coverslips were then coated with 100 µg/ml Rat tail collagen type I (Corning, Corning, NY) dissolved in 0.02 M acetic acid, 1.0 N standardized solution (Alfa Aesar, Wardhill, MA) and kept in the fridge at 4 °C overnight to allow collagen to react with the functionalized surface.

Stiff substrates were generated by coating glass with collagen. Glass coverslips that were 22 mm × 22 mm were treated with hot piranha solution (3:1 H_2_SO_4_ to H_2_O_2_) for 1 h to ensure thorough cleaning and activation. After treatment, the coverslips were immersed in 1% (v/v) 3-aminopropyltriethoxysilane (APTES, Thermo Scientific) solution prepared in 1 mM acetic acid for 2 h to facilitate silanization. The coverslips were then thoroughly rinsed with deionized water and allowed to air dry. Subsequently, the coverslips were baked in an oven at 100°C for 1 h to promote covalent bonding of the APTES layer. Once cooled, the coverslips were treated with 6% (v/v) glutaraldehyde in Dulbecco’s Phosphate-Buffered Saline (DPBS) lacking magnesium and calcium for 2 h to introduce aldehyde functional groups. Following this treatment, the coverslips were extensively washed with deionized water to remove any residual glutaraldehyde. In the next step, glass was coated with 100 µg/ml collagen dissolved in 0.02 M acetic acid and kept in the fridge at 4 °C overnight to allow collagen to adsorb and react with the functionalized glass surface.

### 5.3 Cell seeding and cell sample preparation

After soft (PAA) and stiff (glass) substrates were prepared, cells were allowed to spread onto these substrates in the presence of 10% or 15% FBS in DMEM as the cell line required. Cells were centrifuged 500 times the force of gravity for five min, counted and suspended in culture media at 50,000 # ml^-1^ (MDA-MB-231, HaCaT, WM266-4 and THP-1) or 30,000 # ml^-1^ (HFF) for imaging. Cells were incubated at 37 °C in 5% CO_2_ for 4 h before either fixing and for live cell imaging, cells were incubated for 1h to allow for attachment and they were imaged every 5 min for total duration of 6 h.

### 5.4 Immunofluorescence imaging and analysis

Cells were fixed and stained to visualize their F-actin structure to quantify their response to substrate stiffness through morphological features and assess the F-actin structure. First, the cells were treated with 4% paraformaldehyde (Fisher Scientific) for 10 min, prepared by diluting 16% paraformaldehyde in a cytoskeleton buffer consisting of 10 mM 2-N-morpholino ethane sulfonic acid (MES, pH 6.1, Fisher Scientific), 3 mM MgCl_2_ (Fisher Scientific), 138 mM KCl (Fisher Scientific) and 2 mM EGTA (Sigma Aldrich). Next, cells were permeabilized using a 0.5% Triton-X (Fisher Bioreagents, Waltham, MA) solution in the cytoskeletal buffer for 5 min. Unreacted aldehydes were then blocked with 100 mM glycine (Fisher Scientific) for 15 min. Cells were washed three times with Tris-buffered saline (TBS) for 5 min each. Cells were stained with a 1:400 dilution, Alexa 568-phalloidin (Thermofisher Scientific) for 1 h and 1:500 dilution, DAPI (Sigma) for 15-30 min in TBS containing 0.1% (v/v) Tween-20 (Fisher Bioreagents) and 2% (w/v) bovine serum albumin (BSA, Sigma Life Science). Samples were washed again with TBS three times for 5 min each. To improve image resolution, Prolong Gold mounting media (Thermofisher Scientific) was applied between the substrate with cells and the cover slides. Samples were sealed with VALAP (a mixture of Vaseline, lanolin, and paraffin) and imaged the next day using epifluorescence microscopy with 10x (*NA* =0.3, Nikon, Nishioi, Shinagawa-ku, Japan), 20x (*NA* =0.45, Nikon) and 40x (*NA* = 1.3, Nikon) objectives on a Ti-E microscope (Nikon) with a CMOS (Moment, Photometrics, Tuscan, AZ). Cells were imaged around the edge of the substrate and not the middle as cells were not well-spread in the middle. The edge constitutes the region outside of the center 11 mm x 11 mm region of the 22 mm x 22 mm glass coverslip. The immunofluorescence images were analyzed with ImageJ. Lines around the cell’s edge were drawn and an oval was fit to find the major and minor axis of cell shape. The aspect ratio was calculated as the ratio of the cell’s major axis divided by the cell’s minor axis. The area that the software considered was the shape of the line that fit the cell shape.

Stress fibers were identified using the following image processing approach. The images were processed by using a rolling ball background subtraction method (50 pixels) to enhance the signal of the F-actin stress fibers. Stress fiber length was measured using a free-hand line width of 8 pixels drawn along the F-actin stress fiber. Any line that was 5 μm and longer was considered a stress fiber. The length of each stress fiber and the number of stress fibers was counted. The total stress fiber length was the average stress fiber length multiplied by the average stress fiber per cell. Stress fiber orientation angle was calculated as the difference between the angle of the line that defined the stress fiber and the angle that described the long axis of the cell (Feret angle) with the horizontal axis of the image as a reference angle for each. Thereafter, we categorized these F-actin stress fibers in the cell into three distinct subpopulations as follows: stress fibers less than 30° were marked as parallel, stress fibers between 30° to 60° were marked as transitional and stress fibers greater than 60° were marked as perpendicular. These bins were used to categorize and calculate average stress fiber lengths, numbers of stress fibers per cell and total stress fiber length in a cell in different directions.

### 5.5 Live cell migration imaging and analysis

We fabricated live cell migration chambers by cutting adhesive, heat-resistant, adhesive 1 mm transparent silicone rubber sheets (Laimeisi, Shenzhen, GD, China) using a craft cutter (CriCut Marker 3, Cricut,Inc, South Jordan, UT) to form a chamber on a glass surface. This chamber was filled with clear imaging media, supplemented with 10 or 15% FBS, depending on the cell line and the substrate, with the cells was flipped over onto the chamber. The chamber was sealed with VALAP. The setup was then imaged using phase contrast microscopy using 10x (*NA* =0.3), objectives on a Ti-E microscope with a CMOS, on a heated stage at 37 °C every 5 min for 6 h. This allows for detailed observation and analysis of cell migration behavior. Cells were imaged around the edge of the substrate and not the middle as cells were not well-spread in the middle. The edge constitutes the region outside of the center 11 mm x 11 mm region of the 22 mm x 22 mm glass coverslip. Cell centroids were identified and tracked manually by M - Tracker J plugins in Image J (National Institutes of Health, Bethesda, MD). Single cell speed (*S*) and persistence (*P*) were obtained.

Cell migration was quantified from time-lapse microscopy images acquired at 5 min intervals over 6 h. Analysis was performed only on cells that did not collide with neighbors and remained within the field of view for the entire duration, ensuring complete and unbiased trajectories. We used average instantaneous cell speed in a frame with the interval of 5 min for 6 h. The two-dimensional position vector of the *i*^th^ cell at time, *t*, was denoted as *r_i_* (*t*) = [*x_i_* (*t*), *y_i_* (*t*)]. The instantaneous speed, *S_i_*(*t*), for each interval was calculated using the equation,

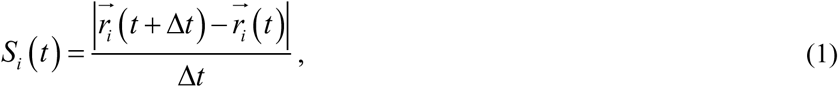

where Δ*t* is 5 min. The mean instantaneous speed, (*S*), for each experimental condition was calculated by averaging *S_i_*(*t*) across all time points and then averaged over all eligible cells. Persistence time, *P*, was calculated from a fit of the mean-squared displacement (MSD) to the persistent random walk model equation,

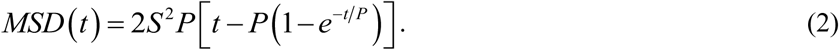

To characterize random cell motility, the mean squared displacement was calculated for individual cell trajectories. Since some MSD data was linear, producing a unique diffusion coefficient (*S*^2^*P*) with compensating values for *S* and *P* that produced similar fits. Consequently, we used instantaneous speed, *S* and the migration random diffusion coefficient, *D,* calculated for a cell from the fit of the MSD and used the following relationship to calculate *P*:

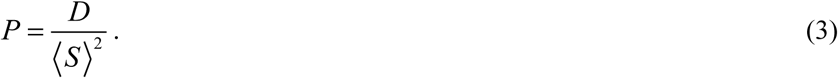

### 5.6 Small interfering RNA (siRNA) transfection

Gene silencing was performed using Silencer® predesigned siRNAs (Thermofisher Scientific) targeting human FHOD3(siRNA, 32380) and DIAPH3 (siRNA, 131029). The corresponding siRNA duplex sequences were as follows: FHOD3: sense 5′-GGAAGUAGCAGAACCACUCtt-3′ antisense,5′-GAGUGGUUCUGCUACUUCCtt-3′. DIAPH3: sense 5′-GCUACAAGCUUUUAAGUCUtt-3′, antisense, 5′-AGACUUAAAAGCUUGUAGCtc-3′. According to the manufacturer’s protocol, oligonucleotides were transfected in cells using Lipofectamine™ RNAiMAX (Thermofisher Scientific). Cells were seeded in a 60 mm culture dish using 2 x 10^6^ cells/5 mL and incubated for 48 h. The complex formed by mixing Lipofectamine™ RNAiMAX and siRNA was added to the cells in the media at siRNA concentrations of 10 μM and 50 μM in MDA-MB-231 and HFF cells, respectively. The control group involved cells transfected with no siRNA and no Lipofectamine RNAiMAX reagent. Cells were incubated for 48 h with the transfection mix, washed with DPBS, and processed for several assays. This included trypsinization for immunofluorescence and live-cell imaging or lysed for RNA transcript analysis.

### 5.7 Total RNA extraction

Total RNA was extracted from cultured cells using the RNeasy Mini Kit (Qiagen, Valencia, CA). For adherent cells, cultures at ∼80% confluence were washed with warm DPBS, detached with 0.25% trypsin-EDTA (Gibco), and pelleted. Cell pellets (2 × 10⁶ adherent and 4 × 10⁶ suspension cells) were lysed in Buffer RLT and homogenized. Following the addition of one volume of 70% ethanol, the mixture was applied to a RNeasy spin column. The column was washed with Buffer RW1 and Buffer RPE according to the manufacturer’s protocol. RNA was eluted in 30 µL of RNase-free water. RNA concentration and purity were determined using a Nanophotometer (Implen Inc, Westlake Village, CA), samples with A₂₆₀/A₂₈₀ ratios between 1.9 and 2.1 were considered suitable for downstream analysis.

### 5.8 Reverse Transcription Polymerase Chain Reaction (RT-PCR)

We used RT-PCR to measure transcript abundance. We used the SuperScript™ III First-Strand Synthesis System (Invitrogen, Carlsbad, CA). For each reaction, 2.5 μg of RNA, with a final concentration of 1 mM of dNTP, 5 ng μl^-1^ of random hexamers were added in the final volume of 30 μL diethylpyrocarbonate treated water. The mix was incubated for 5 min at 65 °C and then 5 min on ice. Following these incubations, 2x RT buffer, 10 mM MgCl_2_, 0.02 M DTT and 4 μg μL^-1^ RNase OUT and superscript III was uniformly mixed and incubated for 50 min at 50 °C, followed by 85 °C for 5 min. Then, 2 uL of RNase H was added to degrade any remnant RNA and incubated at room temperature for 20 min. To prepare the PCR reaction we used Taq buffer, 0.2 mM dNTP mix, 1.5 mM MgCl_2_, 5 U/μL Taq polymerase and a 2.5 μg μL^-1^ DNA template followed by 3 μM for each primer. The PCR reaction program included the following conditions: Initial denaturing temperature was 94 °C for 2 min, denaturing temperature was 94 °C for 1 min, annealing temperature was 55 °C for 30 min, extension temperature was 72 °C for 10 min and 30 cycles were used. Gel electrophoresis through a 2% agarose (60ml, Thermofisher Scientific) was made, to visualize the amplified transcript. PCR amplified products were separated by electrophoresis on 2% agarose gel. We used 1x Prestain (smartGlow prestain, Accuris, Denver, CO) was added to the gel. The 1x loading buffer (Thermofisher Scientific) was added to the amplified products and 100 bp ladder (Thermofisher Scientific, 10488058) prior to loading to the gel. The samples were run at 3.4 V/cm (70 V) for 90 min. Lastly, the gel was imaged and visualized under UV light by a gel imager (smartDoc, Accuris) and data was analyzed using ImageJ. Gene expression was quantified by using ImageJ software. For each band, a region of interest (ROI) was drawn, by selecting a rectangular ROIs around the band and the integrated intensity was measured. This value was corrected by subtracting the intensity of an adjacent local background ROI. The normalized expression was calculated by dividing the background-subtracted intensity of each formin gene by the background-subtracted intensity of the GAPDH loading control from the same sample.

### 5.9 Reverse Transcription Quantitative PCR (RT-qPCR)

We also used RT-qPCR to measure transcript abundance. We used a Luna Universal one-step RT-qPCR kit (New England BioLabs, Ipswich, MA). According to the manufacturer’s protocol, we used, 0.8 μg of total RNA in each reaction. SYBR green master mix, Luna WarmStart RT and 0.4 μM for each primer (Integrated DNA Technologies, Coralville, IA). The RT-qPCR reaction was programmed as follows: Reverse transcription cycle step was at 55 °C for 10 min, initial denaturation step was at 95 °C for 1 min, denaturation step was at 95 °C for 10 s, the extension step was 60 °C for 30 s and the melting curve step temperature was 95 °C and a total of 40 cycles using a Applied Biosystems step OnePlus Real-Time PCR system (Thermofisher Scientific). The primer sequences used in gene profiling included: GAPDH (490 bp), forward (5’-GGGCTCTCCAGAACATCATC-3’), reverse (5’-GACTGAGTGTGGCAGGGACT-3’), DIAPH1 (316 bp), forward (5’–GCT TGT GGC TGA GGA CCT CTC CC-3’), reverse, (5’-GAT CAT AGA CTC AGT CAG AAC AGC-3’),DIAPH2 (324 bp), forward (5’–GGT CAA AGA TTG AAC CCA CAG–3’), reverse (5’-GGT TCT GAA TTA AAG CCT CAC TCA GC-3’), DIAPH3 (307 bp), forward, (5’-GAT CAG ACC TCA TGA AAT GAC TG–3’), reverse, (5’-CTG AAT CAT AGA CTC TGC CAA CCG-3’),FHOD1 (353 bp), forward (5’-CGT GAG CTG AAG CTG GCT GGG GG–3‘), reverse (5’-GCT CTT CCT CCG TGG GCA TCA TGG-3’),FHOD3 (305 bp), forward (5’-CCG ACG CTG CAG AGA ATT CCT GTG G-3’), reverse (5’-GCT TCT CCT CAT CGG TGG GAA T-3’),DAAM1 (319 bp), forward (5’-GCC CGA GAA CAA ACT GGA AGG-3’), reverse (5’-GGG CAG ATC TTC CTG TTC GTC C-3’),DAAM2 (296 bp), forward (5’-GGA GCG TGT CCC TGG CAC CGT ATG G-3’), reverse (5’-CCA GCA TGT CCT TAG CAA GGT CCT CC-3’),FMN1 (179 bp), forward (5’–GCT GAA GAA GGG GGC TAC CGC–3’), reverse (5’–GGA GAG TGG GAG TGG CCT TCG–3’),FMN2 (356 bp), forward (5’-GCC TCT TTA CTG GAC CAG G-3’), reverse (5’-CGA GTT CGTCTG ACT GTG C-3’),FMNL1 (301 bp), forward (5’–GCA CTG AAA CCC AGC CAG ATC ACC–3’), reverse (5’–GGA AGT CCA GGC CCA GAG CCT GC–3’),FMNL2 (326 bp), forward (5’-GCT CTG AAG CCC AAT CAG ATC AAT GGC-3’), reverse (5’-GGT AGG AAC CGC ATC AAG CAT TCC-3’),FMNL3 (318 bp), forward (5’-GCA CTG AAA CCC AAC CAG ATC AGT GGC-3’), reverse (5’-GCG CAT CAG GCA CTC CAC GAA GTC C-3’),INF1 (280 bp), forward (5’-GGA CCT TGG CAG CCA GGC AGG-3’), reverse (5’-CGC AAG GTC TCT GAT CCA TAA TGC-3’), INF2 (338 bp), forward (5’-GCT GCC ATC CAA CGT GGC ACG TGA GC-3’), reverse (5’-CGT GCT TCT CGG GAA GGA GCT TAA GG-3’), DELP (delphilin) (289 bp), forward (5’-GGC ACC ATC TGG GGT CAG CTC GGG G-3’), reverse (5’-GCA GCA GCT GCG CGA GAT GTG CGG GC-3’).

To analyze the data collected from Applied Biosystems step OnePlus Real-Time PCR system by using step one plus software to calculate the threshold cycles (*Ct*). We used the 2^-*ΔΔCt*^ a method of relative quantification that is most frequently found in popular software packages for qPCR experiments. This method used *Ct* information generated from a qPCR system to calculate relative formin gene expression. To correct differences in the amount of RNA added for each sample and to reduce variation caused by PCR set-up and the cycling process, reference genes or internal control genes have been used to normalize the PCRs. Housekeeping genes (GAPDH) were used as reference genes because their expression levels remain relatively stable in response to any treatment. The following equation represents a measure that is proportional to expression:

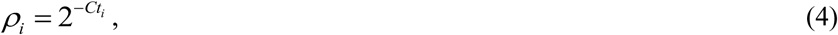

where *i* is the specific gene transcript. Normalized expression, *ϕ_i_*, is defined as follows:

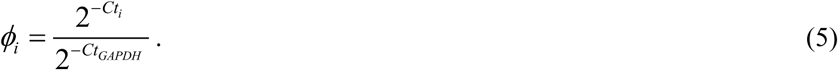

The normalized concentration final result of this method is presented as the fold change of target gene expression in a target sample relative to GAPDH. Heat maps of normalized expression was generated in excel, the centered color scale (yellow-orange-red) shows upregulation (red) and down regulation (yellow) relative to the average.

## Abbreviations

DPBS: Phosphate-Buffered Saline
RLT: RNA Lysis Tissue buffer
BME: (β-MercaptoEthanol)
APS: Ammonium Persulfate
APTES: 3-AminoPropylTriEthoxySilane
EGTA: Ethylene Glycol bis(2-aminoethyl ether)-N,N,N′,N′-Tetraacetic Acid
BSA: Bovine Serum Albumin
ECM: ExtraCellular Matrix
TEMED: N,N,N’, N’-Tetraacetylethylenediamines
TBS: Tris Buffered Saline
HEPES: 2-HydroxyEthylPiperazine-1-Ethane Sulfonic acid
FBS: Fetal Bovine Serum
MES: 2-N-Morpholino Ethane Sulfonic acid
HFF: Human Foreskin Fibroblast
PAA: PolyAcrylAmide
GAPDH: Glyceraldehyde 3-Phosphate Dehydrogenase
FHOD3: Formin Homology 2 Domin Containing 3
DIAPH3: Diaphanous Related Formin 3
KD: Knock-Down
FG: Functionalized Glass
0.2: 200 Pa PAA gel
2: 2000 Pa PAA gel
20: 20,000 Pa PAA gel

## Author Contributions

FRN: Conceived experiments, conducted experiments, analyzed and interpreted the data across all figures, prepared the figures, wrote the article and edited the article. AF: Prepared soft and stiff substrates used throughout the experiments, wrote the article and edited the article. ICS: Conceived experiments, analyzed and interpreted the data across all figures, prepared the figures, wrote the article and edited the article.

## Acknowledgements

We acknowledge the DNA Facility and David Wright at Iowa State University. We acknowledge the Nigel Reuel Group for use of the Nanophotometer and the PCR Touch Thermal Cycler.

This work was supported by the National Institutes of Health under grant No. GM143302.

## Data Availability Statement

The data that support the findings of this study are available from the corresponding author upon reasonable request.

