## Supplemental Figures for "FHOD3 and DIAPH3 control cell migration and differentially shift the balance of parallel and perpendicular stress fibers"

### FHOD3 and DIAPH3 control stress fibers and cell migration through shared and distinct mechanisms

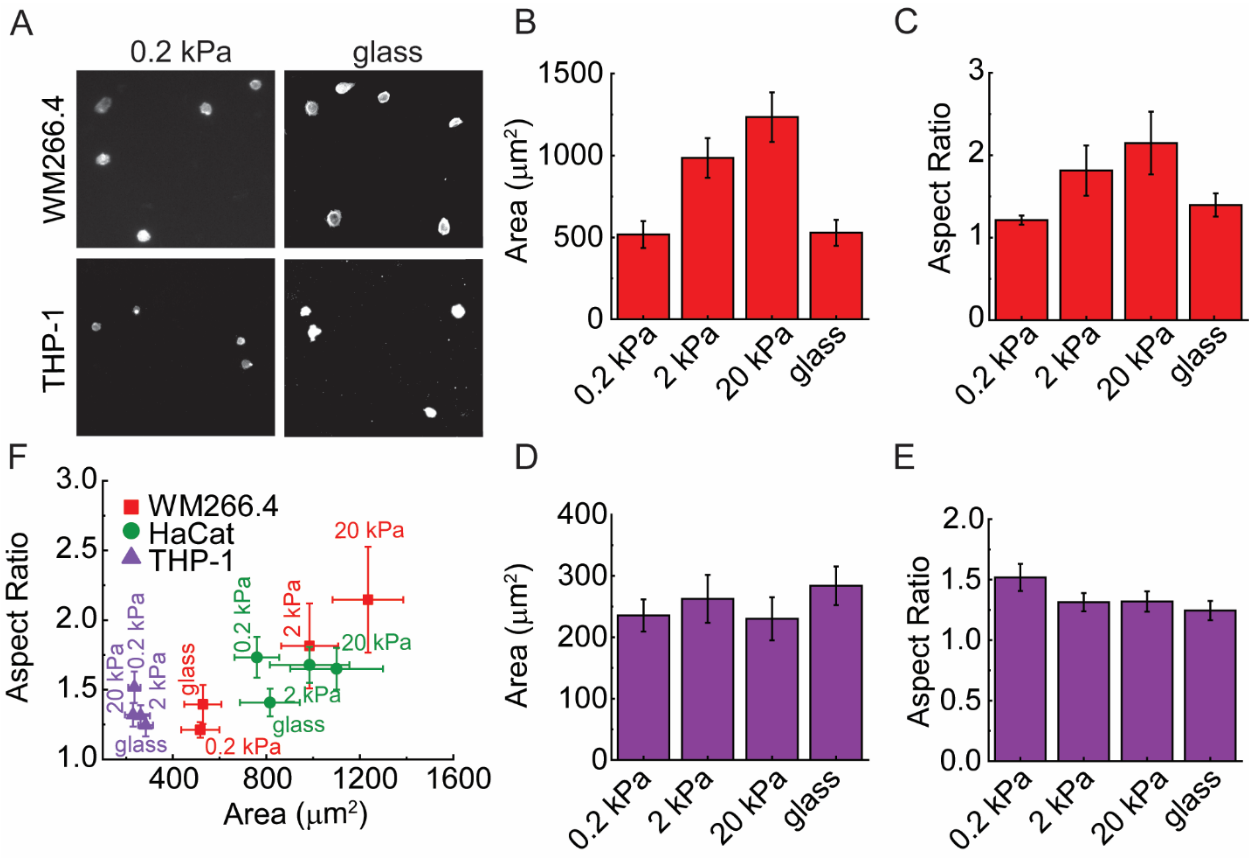


**Supplementary Figure 1: Cells with different migration modes show different morphological and behavioral responses on substrate stiffness.** (A) Scatter plot of cell aspect ratio versus cell area for cell types cultured on collagen-coated on 0.2,2,20 kPa PAA and glass substrates. This analysis reveals distinct morphological niches occupied by each cell type in response to mechanical cues. (B) Representative immunofluorescence images of F-actin (phalloidin-568) in WM266.4 and THP-1 cells, illustrating their specific morphological adaptations to 0.2 kPa PAA and glass microenvironments. Scale bar: 100 µm. (C-F) Systematic quantification of cell area (C, E) and aspect ratio (D, F) for WM266.4 and THP-1 cells across different substrate stiffness, showing their unique mechanoresponsive profiles. $N_{samples}$ =3, $N_{cells,\ HFF}$ = 60, and $N_{cells,\ MDA\text{-}MB\text{-}231}$ = 60. $N_{cells,\ HaCat}$ = 60. $N_{cells,\ WM266\text{-}4}$ = 60. $N_{cells,\ THP\text{-}1}$ = 60. Error bar represents a 95% confidence interval unless otherwise stated. All the experiments were replicated at least four times unless otherwise stated.

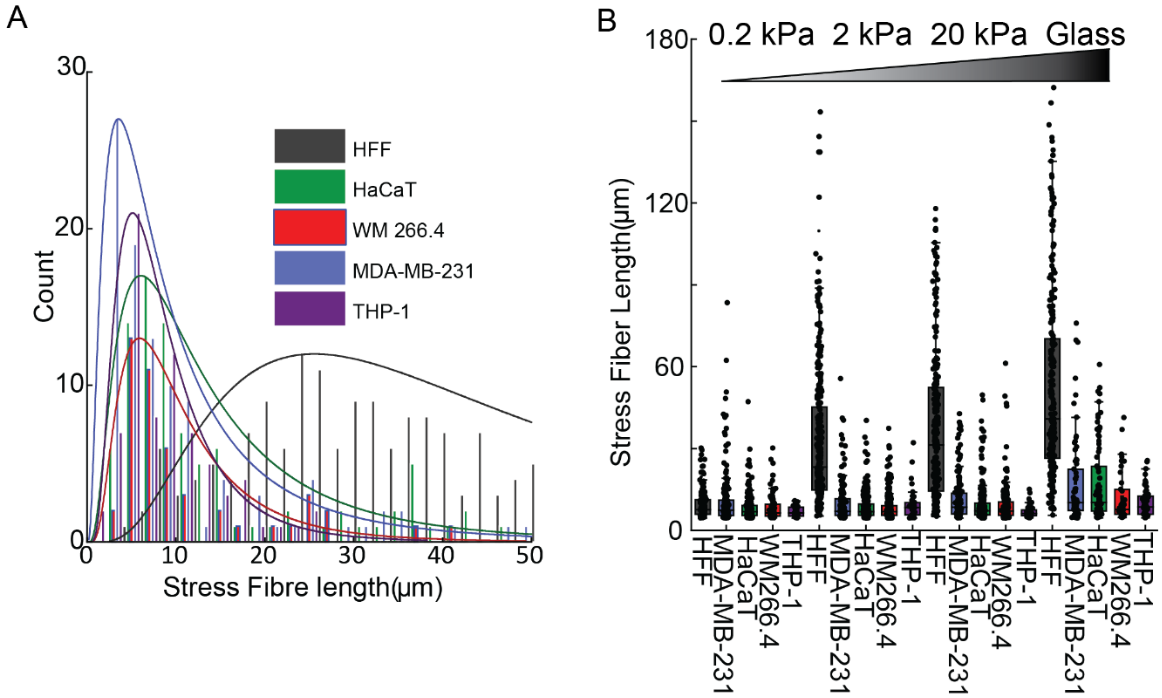


**Supplementary Figure 2: Stiffness regulates actin fiber architecture across different cell types and migration modes.** (A) Quantification of mean stress fiber length in HFF, MDA-MB-231s, and HaCaT WM266.4 and THP-1 cells plated on 0.2 kPa PAA and glass substrates. Indicating that only fibers ≥5 μm in length were considered as stress fibers in the analysis. (B) The proportion of analyzed fibers that met the ≥5 μm length threshold across all cell types and substrates. $N_{cells,\ HFF}$ = 10, and $N_{cells,\ MDA\text{-}MB\text{-}231}$ = 10, 37 < $N_{Stress\ Fibres}$, < 333, and $N_{cells}$ =10. All the experiments were replicated at least three times unless otherwise stated.

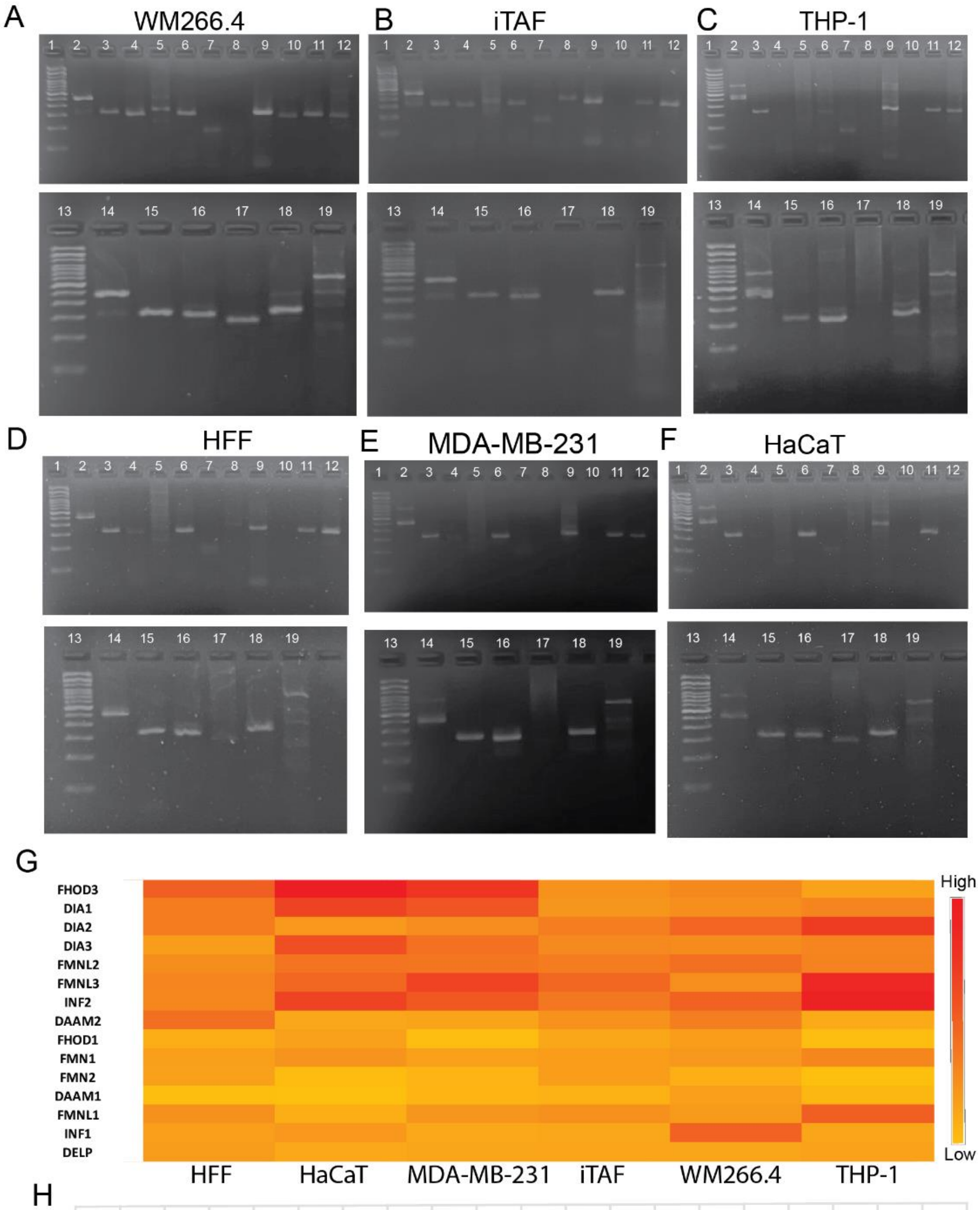


H

| LANE | NAME | BP SIZE |
|---|---|---|
| 1 | LADDER (100)bp | |
| 2 | GAPDH | 490bp |
| 3 | DIA1 | 316bp |
| 4 | DAAM2 | 296bp |
| 5 | FHOD1 | 343bp |
| 6 | FHOD3 | 305bp |
| 7 | FMN1 | 179bp |
| 8 | FMN2 | 356bp |
| 9 | DIA2 | 307bp |
| 10 | DAAM1 | 319bp |
| 11 | DIA3 | 324bp |
| 12 | FMNL1 | 301bp |
| 13 | LADDER (100)bp | |
| 14 | GAPDH | 490bp |
| 15 | FMNL2 | 327bp |
| 16 | FMNL3 | 327bp |
| 17 | INF1 | 280bp |
| 18 | INF2 | 338bp |
| 19 | DELPHILLIN | 301bp |

**Supplementary Figure 3: Expression profiling of formin isoforms across mammalian cell lines Showing qualitative analysis quantified by gel express method.** (A-F) Reverse transcription -PCR (RT-PCR) analysis of fifteen formin isoforms and GAPDH (Loading control) in HFF, MDA-MB-231, HaCaT, WM266-4, THP-1, and iTAF cells. Inverted grayscale images show agarose gel electrophoresis of the PCR product after 40 amplification cycles. (G) Heat map of formin expression levels quantified from band intensities normalized to GAPDH across the six cell lines. Color scale: Yellow (low) to red (High). (H) Table showing the expected PCR product band sizes for formin PCR products, on color images of agarose gel electrophoresis.  (G) A heat map showing expression of formin across six cell lines. (H) Representation of names, of formin PCR products and their base size on agarose gel Images. Experimental design: n=2 biological replicates and n=2 technical replicates.

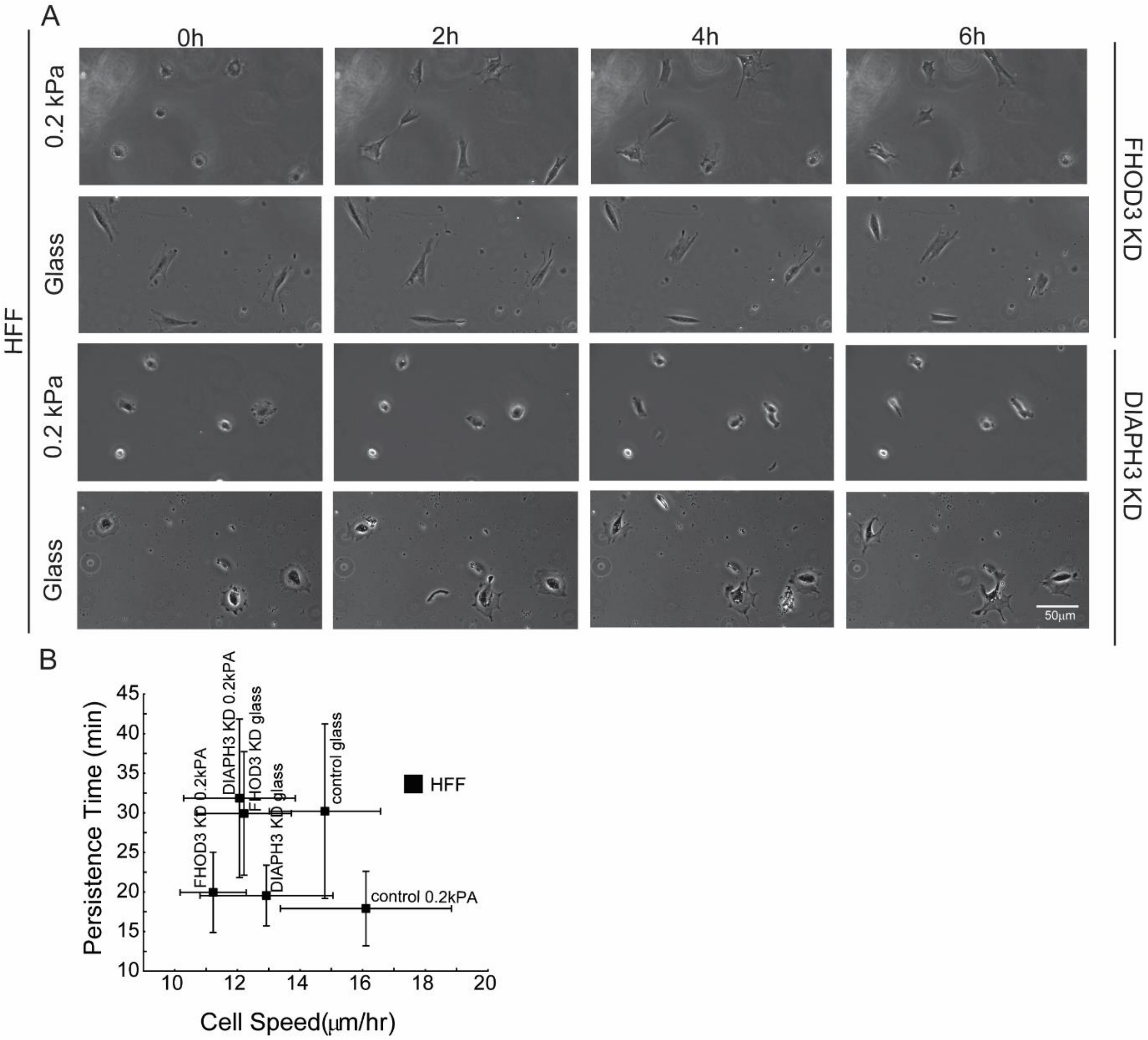


**Supplementary Figure 4: FHOD3 and DIAPH3 regulate cell morphology and migration dynamics in HFF cells.** (A) A phase contrast time-lapse montage of MDA-MB-231 cells migrating on 0.2kPa and glass substrates following transfection with *FHOD3* or *DIAPH3* targeting siRNA. Scale bar:100μm. (B) Scatter plot correlating cell speed with persistence time (the duration of movement before turning) for cells on 0.2kPa and glass substrates before and after *FHOD3* or *DIAPH3* knockdown.$N_{cells,\ HFF} > 55$, and $N_{cells,\ MDA\text{-}MB\text{-}231} > 55$. Error bar represents a 95% confidence interval unless otherwise stated. P-values were evaluated by using a two-tailed unpaired Student *t*-test. Significant * represents $p < 0.005$. All the experiments were replicated at least three times unless otherwise stated.

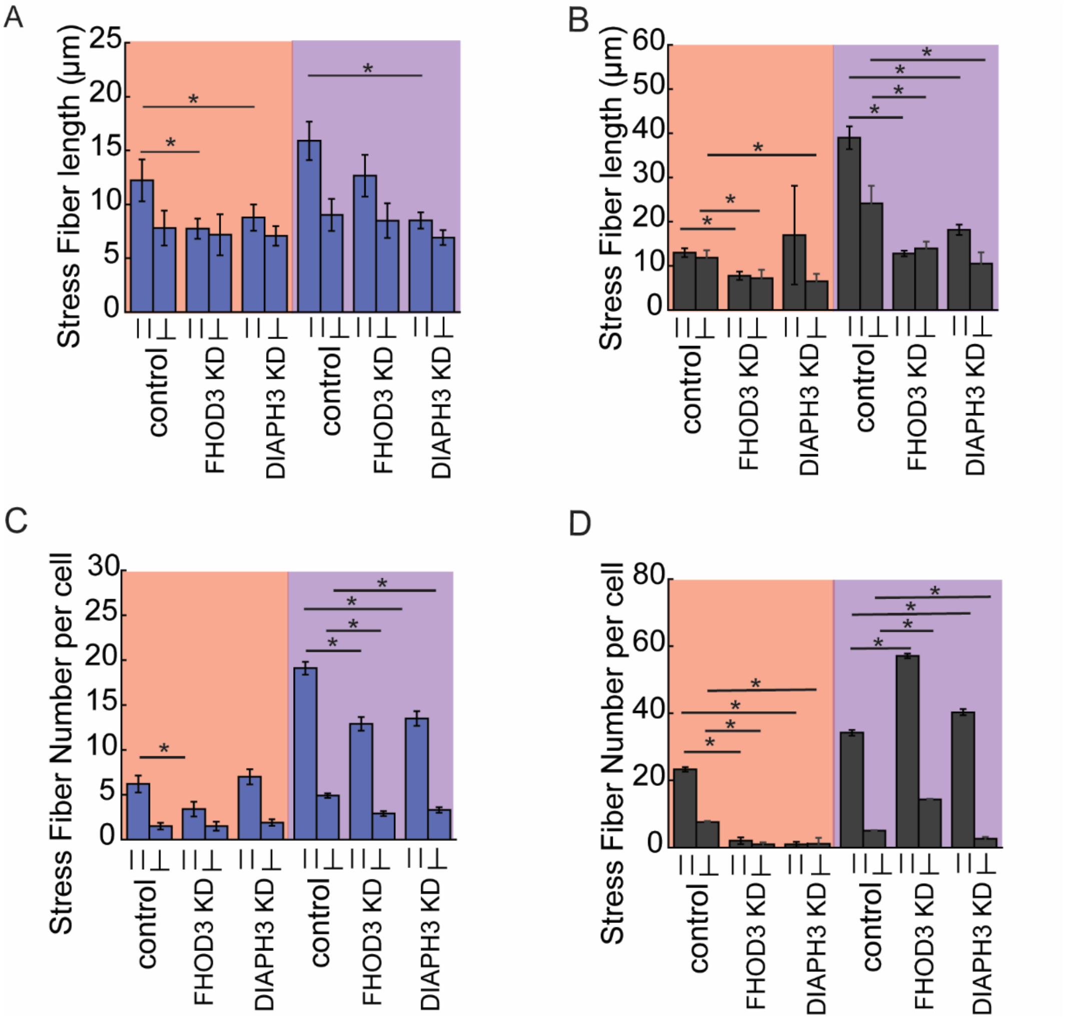


**Supplementary Figure 5: FHOD3 and DIAPH3 regulate distinct stress fibers in MDAMB-231s and HFF cells.** (A, B) Quantification of parallel and perpendicular stress fiber length in (A) MDA-MB-231 and (B) HFF cells following FHOD3 or DIAPH3 knockdown on 0.2 kPa PAA and glass substrates. (C, D) Quantification of the number of parallel and perpendicular stress fibers per cell in (C) MDA-MB-231 and (D) HFF cells under the same conditions $N_{cells, HFF}$ =10, and $N_{cells, MDA\text{-}MB\text{-}231}$ >10, 25 < $N_{Stress\ Fibers}$ < 201, Error bars represent a 95% confidence interval unless otherwise stated. P-values were calculated using a two-tailed unpaired Student's *t*-test. (Significant *represents $p < 0.005$. All the experiments were replicated at least three times unless otherwise stated.
